# Evaluating Autonomous LLM Agents Across Molecular Prediction and Optimization Benchmarks

**DOI:** 10.64898/2026.10.01.755314

**Authors:** Khachik Smbatyan, Tsolak Ghukasyan, Garik Petrosyan

## Abstract

Large language model (LLM) agents are increasingly capable of carrying out autonomous computational research, but it remains unclear whether they can develop molecular modeling methods that compete with strong human-developed approaches. Here, we evaluate autonomous method development across four settings: Therapeutics Data Commons (TDC) ADMET tasks, the OpenADMET ExpansionRx Challenge, the activity prediction track of the OpenADMET PXR Induction Challenge, and the Practical Molecular Optimization (PMO) benchmark. Agents followed the prescribed data splits, metrics, and evaluation protocols for each benchmark. On TDC, the best agent-developed score across configurations improved on the leaderboard reference on 18 of 21 tasks and on the combined leaderboard and peer-reviewed reference on 17, while individual Codex configurations improved on the combined reference on 13 tasks. On ExpansionRx, five-agent and single-agent Codex configurations achieved overall scores corresponding to fourth and sixth place relative to the published final leaderboard, with the five-agent configuration reaching top-five performance on seven of nine endpoints. On PXR, five-agent Codex achieved a mean absolute error slightly outperforming the best published challenge entry. On PMO, five-agent Codex developed a molecular optimizer that outperformed existing methods evaluated under the benchmark’s standard setup; other methods obtained higher reported scores under modified conditions, including additional molecular pretraining data or a different oracle-call budget. Together, these results show that current LLM agents can autonomously develop high-performing molecular prediction and optimization methods across substantially different drug-discovery settings, in several cases matching or exceeding leading human-developed approaches.

## 1. Introduction

Predicting molecular absorption, distribution, metabolism, excretion, and toxicity (ADMET) is an important component of drug discovery, helping researchers prioritize compounds for experimental testing and identify potential liabilities during optimization. Machine learning models can support these decisions by estimating molecular properties from chemical structures, enabling the evaluation of large compound collections [1]. Developing effective predictors involves decisions about data preparation, molecular representations, model architectures, training procedures, and validation strategies. These choices must account for differences in dataset size, experimental origin, and the property being predicted. ADMET model development therefore provides a practical setting for studying the automation of computational research, combining a scientifically relevant objective with an iterative process of method design and evaluation. Beyond property prediction, molecular optimization requires methods that propose and search for molecules with desirable properties under limited scoring budgets. Prediction and optimization therefore provide complementary settings for assessing autonomous method development in drug discovery.

Large language models (LLMs) equipped with coding tools and execution environments can perform several steps in this process, including implementing candidate methods, running experiments, interpreting results, and revising their approaches. Evaluations of such agents have demonstrated their ability to conduct machine learning experiments, while also revealing substantial variation in success across tasks and runs [2, 3]. Their usefulness for molecular modeling depends on whether these capabilities translate into competitive prediction and optimization methods and reproducible workflows within available computational resources. Evaluating this potential requires considering both the quality of the resulting models and the organization of the agents developing them, including the underlying LLM, the software environment through which it operates, and the number of collaborating agents.

Recent work has explored LLM-based agents for autonomous development of molecular prediction models, including tasks from the Therapeutics Data Commons (TDC). DrugAgent combines planning and coding agents to propose, implement, and evaluate modeling approaches, with experiments on PAMPA, HIV, and DAVIS datasets [4]. MolAgent integrates agents with the AutoMol backend to automate data preparation, feature selection, model training, and ensembling through expert-designed workflows [5]. Closed-loop Auto Research evaluates agent-driven changes to molecular representations, model code and external training data across 22 TDC ADMET endpoints. Its validation-selected pipeline outperformed the starting baseline on 12 of them, with a mean normalized held-out improvement of 0.013; however, substantial validation gains did not always transfer to the test set [6]. Whereas these approaches incorporate curated domain guidance, predefined modeling backends, or restrictions on pipeline modifications, we set out to evaluate end-to-end method development while leaving modeling choices to the agents, subject to the task and resource constraints.

In this work, we evaluate autonomous method development on 21 tasks from the TDC ADMET Benchmark Group [7], comparing the resulting predictors with reference scores from the TDC leaderboards and peer-reviewed publications. We compare performance across harness–model pairings and agent counts and describe the modeling strategies developed by the agents. We extend this evaluation to the nine regression endpoints of the OpenADMET ExpansionRx Challenge [8], which tests predictions for later-stage compounds using earlier lead-optimization data. We additionally evaluate agent capabilities on the OpenADMET PXR Induction Challenge dataset, a focused prediction problem involving chemical analogues of screening hits [9, 10]. Finally, we evaluate the agents on the 23-task Practical Molecular Optimization (PMO) benchmark, which measures the discovery of high-scoring molecules under a limited oracle-call budget [11]. PMO extends the evaluation to autonomous development of molecular optimizers, requiring candidate generation and search strategies that use scoring evaluations efficiently. Although both TDC and ExpansionRx evaluate ADMET prediction, they represent distinct modeling settings, with different properties, chemical contexts, and evaluation and feedback regimes. We therefore treat them as complementary benchmarks rather than interchangeable evaluations of the same task. Together, these benchmarks span molecular property prediction under scaffold-based, temporal, and within-series evaluation settings, as well as molecular optimization.

## 2. Results

### 2.1. Performance on the TDC ADMET Benchmark

The TDC ADMET Benchmark Group comprises 22 molecular property prediction tasks spanning absorption, distribution, metabolism, excretion, and toxicity, with predefined scaffold-based splits and task-specific evaluation metrics. We evaluated the ability of LLM agents to develop predictive models for 21 tasks from the TDC ADMET Benchmark Group, spanning regression and classification endpoints. Within each run, agents were required to develop a single method applicable to all tasks rather than an independent method for every task. Experiments compared two harness–model pairings — Codex with GPT-6 Astra and Claude Code with Opus 5 — at agent counts of 1, 5, and 10, together with a mixed configuration of five Claude Code–Opus 5 agents and five Codex–GPT-6 Astra agents, under the resource limits described in Methods.

Reference scores for each task are drawn from two cumulative sets: TDC leaderboard entries (denoted TDC in Table 1) [12], and leaderboard entries combined with peer-reviewed publications (denoted TDC+PR), which improves the reference on 15 of the 21 tasks. Taking the best score per task across configurations, agent-developed models improved on the leaderboard reference on 18 tasks and on the combined reference on 17 (Table 1), while single-agent Codex and five-agent Codex each improved on the combined reference on 13 tasks despite using one method across all 21. Notable improvements across configurations include plasma protein binding (MAE 7.229 to 6.431), lipophilicity (MAE 0.393 to 0.359), and half-life (Spearman correlation 0.619 to 0.672). Increasing agent count does not consistently improve performance: five-agent Codex outperforms ten-agent Codex on 13 tasks, with two ties, while single-agent Claude Code outperforms its five- and ten-agent counterparts on 17 and 15 tasks, respectively. The mixed configuration improves on the combined reference on six tasks, compared with eight for ten-agent Codex, and achieves no best score among the evaluated agentic configurations.

**Table 1.** Reference and agentic performance on the TDC ADMET benchmark group. Results cover 21 endpoints. TDC denotes the best website score; TDC+PR combines website leaderboard entries with peer-reviewed publications (specified in square brackets). Bold endpoint scores indicate improvements over TDC+PR. Best agentic is the best score per endpoint across the seven named configurations. The final rows count strictly better scores using the displayed values; ties are excluded.

| Endpoint | Metric | TDC | TDC+PR | CC-1 | CC-5 | CC-10 | CX-1 | CX-5 | CX-10 | Mixed-10 | Best agentic |
| --- | --- | --- | --- | --- | --- | --- | --- | --- | --- | --- | --- |
| Caco-2 | MAE ↓ | 0.2560 | 0.2560 | 0.3200 | 0.2790 | 0.2740 | 0.2660 | 0.2720 | 0.2720 | 0.2850 | 0.2660 |
| HIA | AUROC ↑ | 0.9930 | 0.9938<sup>[54]</sup> | **0.9940** | 0.9850 | 0.9870 | **0.9940** | 0.9880 | 0.9860 | 0.9830 | **0.9940** |
| Pgp | AUROC ↑ | 0.9380 | 0.9430<sup>[55]</sup> | **0.9520** | 0.9360 | 0.9370 | **0.9470** | **0.9440** | 0.9420 | **0.9460** | **0.9520** |
| Lipophilicity | MAE ↓ | 0.4560 | 0.3930<sup>[56]</sup> | 0.4560 | 0.4850 | 0.4570 | **0.3590** | 0.3940 | 0.4070 | 0.3950 | **0.3590** |
| Solubility | MAE ↓ | 0.7410 | 0.7140<sup>[50]</sup> | 0.7200 | 0.7230 | 0.7380 | **0.6690** | **0.6850** | **0.7120** | **0.6990** | **0.6690** |
| BBB | AUROC ↑ | 0.9240 | 0.9240 | 0.9190 | 0.9140 | **0.9260** | 0.9240 | **0.9260** | 0.9220 | 0.9120 | **0.9260** |
| PPBR | MAE ↓ | 7.4400 | 7.2290<sup>[57]</sup> | **6.7860** | **7.0870** | **7.1280** | **6.4310** | **6.6300** | **6.7480** | **6.5920** | **6.4310** |
| VDss | rho ↑ | 0.7130 | 0.7160<sup>[54]</sup> | **0.7170** | 0.7130 | 0.6990 | **0.7240** | **0.7280** | 0.7150 | 0.7070 | **0.7280** |
| CYP2C9 inhibition | AUPRC ↑ | 0.8590 | 0.8770<sup>[55]</sup> | 0.8140 | 0.8100 | 0.7740 | **0.8930** | **0.8930** | **0.8900** | **0.8870** | **0.8930** |
| CYP2D6 inhibition | AUPRC ↑ | 0.7900 | 0.8110<sup>[55]</sup> | 0.7220 | 0.7310 | 0.7180 | **0.8320** | **0.8200** | **0.8240** | **0.8270** | **0.8320** |
| CYP3A4 inhibition | AUPRC ↑ | 0.9160 | 0.9230<sup>[55]</sup> | 0.8990 | 0.8930 | 0.8810 | **0.9350** | **0.9350** | **0.9360** | **0.9340** | **0.9360** |
| CYP2C9 substrate | AUPRC ↑ | 0.4740 | 0.4740 | 0.4000 | 0.3760 | 0.4140 | 0.4650 | 0.4690 | 0.4660 | 0.4470 | 0.4690 |
| CYP2D6 substrate | AUPRC ↑ | 0.7360 | 0.7660<sup>[55]</sup> | 0.7120 | 0.6990 | **0.7670** | 0.7350 | 0.7030 | 0.7120 | 0.7070 | **0.7670** |
| CYP3A4 substrate | AUROC ↑ | 0.6670 | 0.6920<sup>[57]</sup> | 0.6620 | 0.6570 | 0.6700 | 0.6740 | 0.6410 | 0.6410 | 0.6540 | 0.6740 |
| Half-life | rho ↑ | 0.5760 | 0.6189<sup>[58]</sup> | **0.6260** | 0.5810 | 0.5800 | **0.6720** | **0.6300** | **0.6200** | 0.5990 | **0.6720** |
| Hepatocyte clearance | rho ↑ | 0.5360 | 0.5360 | 0.4970 | 0.4850 | 0.4630 | **0.5450** | **0.5510** | 0.5100 | 0.4990 | **0.5510** |
| Microsome clearance | rho ↑ | 0.6300 | 0.6370<sup>[50]</sup> | 0.6270 | 0.6170 | 0.5840 | **0.6520** | **0.6570** | **0.6420** | 0.6260 | **0.6570** |
| LD50 | MAE ↓ | 0.5520 | 0.5380<sup>[56]</sup> | **0.5340** | 0.5700 | 0.5910 | 0.5380 | **0.5370** | **0.5360** | 0.5430 | **0.5340** |
| hERG | AUROC ↑ | 0.8800 | 0.8800 | 0.8470 | 0.8250 | **0.8880** | 0.8490 | 0.8400 | 0.8580 | 0.8640 | **0.8880** |
| AMES | AUROC ↑ | 0.8710 | 0.8720<sup>[54]</sup> | **0.8750** | **0.8760** | 0.8520 | **0.8730** | **0.8730** | 0.8700 | 0.8660 | **0.8760** |
| DILI | AUROC ↑ | 0.9560 | 0.9560 | 0.9200 | 0.9260 | 0.9030 | 0.9230 | 0.9030 | 0.9240 | 0.9340 | 0.9340 |
| **Wins vs TDC** | — | — | — | **8/21** | **4/21** | **7/21** | **15/21** | **14/21** | **11/21** | **9/21** | **18/21** |
| **Wins vs TDC+PR** | — | — | — | **7/21** | **2/21** | **4/21** | **13/21** | **13/21** | **8/21** | **6/21** | **17/21** |

Configurations: CC, Claude Code with Opus 5; CX, Codex with GPT-6 Astra (max); Mixed-10, five CC-Opus 5 agents plus five CX-GPT-6 Astra (max) agents. Numeric suffixes give agent counts.
Metrics: MAE, mean absolute error; AUROC, area under the receiver operating characteristic curve; AUPRC, area under the precision-recall curve; rho, Spearman rank correlation. Downward arrows indicate lower is better; upward arrows indicate higher is better.
Endpoints: HIA, human intestinal absorption; Pgp, P-glycoprotein inhibition; BBB, blood-brain barrier penetration; PPBR, plasma protein binding; VDss, volume of distribution at steady state; LD50, acute toxicity; DILI, drug-induced liver injury.

The top-performing agent-developed approaches were developed by single- and five-agent Codex running GPT-6 Astra. They used endpoint-specific TabM [13] ensembles built from selected combinations of frozen pretrained molecular representations (3D conformation-based Uni-Mol2 embeddings [14] and 2D molecular graph-based MiniMol embeddings [15]), fingerprints, and chemical and ionization descriptors. A more detailed description is provided in Supplementary Text S1.1.

To assess whether autonomous method development extends beyond TDC, we also evaluate the OpenADMET ExpansionRx Challenge, the activity prediction track of the OpenADMET PXR Induction Challenge, and the PMO benchmark. These settings test prediction during lead optimization, activity prediction within chemical series, and the discovery of high-scoring molecules under a limited oracle-call budget, respectively.

### 2.2. Evaluation on the OpenADMET ExpansionRx Challenge

The OpenADMET ExpansionRx Challenge [8] evaluates nine ADMET regression endpoints using data from drug discovery campaigns at Expansion Therapeutics. The tasks cover lipophilicity, kinetic solubility, human and mouse microsomal intrinsic clearance, Caco-2 permeability and efflux, and unbound fractions in mouse plasma, brain, and muscle. Training compounds come from earlier stages of lead optimization, while test compounds come from later stages of the same campaigns. This temporal split complements the scaffold-based TDC evaluation by assessing predictions for compounds encountered as optimization progresses.

The challenge ranks individual endpoints by mean absolute error (MAE) and overall performance by macro-averaged relative absolute error (MA-RAE), with lower values indicating better performance. We replayed the challenge within 24 hours, retaining its official training and test sets and feedback stages, with a cap of 30 submissions compared with approximately 85 opportunities under the original schedule. The original half-test feedback membership was unavailable, so the replay used a fixed random half of the test compounds. Table 2 compares the best reported score for each endpoint with the single-agent and five-agent Codex configurations used for TDC. The overall reference comes from the FINAL leaderboard evaluated on the full test set. Its leading entry, pebble, achieved an MA-RAE of 0.511 ± 0.007. This overall result belongs to one complete submission; the endpoint-specific best scores may come from different submissions [16].

**Table 2.** Reference and agentic performance on the OpenADMET ExpansionRx Challenge. Rows report the nine endpoint MAEs and overall MA-RAE from the full test set. All metrics are lower-is-better. Best challenge reports the central score displayed by the official leaderboard. Ranks in parentheses give the position the configuration would occupy relative to the published final leaderboard, followed by the number of ranked entries (N).

| Endpoint | Metric | Best challenge | CX-1 (rank/N) | CX-5 (rank/N) | Best agentic (rank/N) |
| --- | --- | --- | --- | --- | --- |
| LogD | MAE ↓ | 0.2820 | 0.3202 (8/104) | 0.3129 (4/104) | 0.3129 (4/104) |
| KSOL | MAE ↓ | 0.2708 | 0.3039 (13/104) | 0.2924 (7/104) | 0.2924 (7/104) |
| MLM CLint | MAE ↓ | 0.3054 | 0.3237 (4/104) | 0.3230 (4/104) | 0.3230 (4/104) |
| HLM CLint | MAE ↓ | 0.2485 | 0.2577 (7/104) | 0.2508 (4/104) | 0.2508 (4/104) |
| Caco-2 efflux | MAE ↓ | 0.2723 | 0.2990 (5/104) | 0.2987 (5/104) | 0.2987 (5/104) |
| Caco-2 Papp A>B | MAE ↓ | 0.1911 | 0.2022 (3/104) | 0.2119 (3/104) | 0.2022 (3/104) |
| MPPB | MAE ↓ | 0.1451 | 0.1905 (21/104) | 0.1762 (15/104) | 0.1762 (15/104) |
| MBPB | MAE ↓ | 0.1097 | 0.1308 (13/104) | 0.1200 (3/104) | 0.1200 (3/104) |
| MGMB | MAE ↓ | 0.1410 | 0.1720 (39/104) | 0.1427 (2/104) | 0.1427 (2/104) |
| **Overall (FINAL)** | **MA-RAE ↓** | **0.5113** | 0.5574 (6/104) | 0.5378 (4/104) | 0.5378 (4/104) |

Best agentic is the lowest score for each endpoint across the configurations. In the overall row, it is the lowest MA-RAE achieved by a single complete configuration, rather than an aggregate formed by selecting different configurations for different endpoints. Reference values retain the leaderboard precision; bootstrap uncertainties are omitted from the table for compactness. Snapshot: 22 September 2026.
Configurations follow Table 1: CX, Codex with GPT-6 Astra (max). Numeric suffixes indicate agent counts.
Endpoints: LogD, logarithm of the octanol–water distribution coefficient; KSOL, kinetic solubility; MLM/HLM CLint, mouse/human liver microsomal intrinsic clearance; Papp A>B, apical-to-basolateral apparent permeability; MPPB, mouse plasma protein binding; MBPB, mouse brain protein binding; MGMB, mouse gastrocnemius muscle binding. Binding endpoints are reported as percent unbound. MAE uses LogD directly and the challenge logarithmic transformation for the other endpoints (Section 5.1).

Throughout these challenge comparisons, reported leaderboard positions indicate where an agentic submission would fall in the published leaderboard ordering, but do not imply statistically significant separation from neighboring entries.

Five-agent Codex achieved an overall MA-RAE of 0.5378 ± 0.0068, corresponding to fourth place relative to the final leaderboard and 5.2% higher error than the leading score of 0.5113 (Table 2). The uncertainty is the standard deviation of the organizer-style bootstrap distribution. Single-agent Codex achieved an MA-RAE of 0.5574, corresponding to sixth place. These placements compare each configuration independently with the published challenge entries. The five-agent configuration reduced overall MA-RAE by 3.5% relative to the single agent in these runs; this comparison does not estimate variation across repeated agent development runs.

Strong performance extended across individual endpoints: five-agent Codex attained scores corresponding to top-five placements on seven of the nine endpoints, including three in the top three. Its MGMB MAE of 0.143 would place second and was only 1.2% higher than the leading score of 0.1410. For HLM CLint, its result would place fourth with an MAE of 0.251, only 0.9% higher than the leading score of 0.249. Five-agent Codex would also place third for MBPB and Caco-2 Papp (A>B), fourth for LogD and MLM CLint, and fifth for Caco-2 efflux. Across configurations, the five-agent configuration achieved lower MAE on eight of nine endpoints, with its largest relative improvement over the single agent on MGMB (17.0%). Single-agent Codex performed better on Caco-2 Papp A>B (0.202 versus 0.212), with both scores corresponding independently to third place.

The solution developed by five-agent Codex combined graph neural networks, multitask tabular models (including TabM [13]), fine-tuned molecular representation models (MiniMol, Uni-Mol [17]) and kernel predictors in endpoint-specific ensembles, supported by public-assay pretraining and calibration (detailed description in Supplementary Text S1.2).

### 2.3. Evaluation on the OpenADMET PXR Induction Challenge

The OpenADMET PXR Induction Challenge [9] provides a complementary setting for evaluating autonomous molecular model development. Its activity prediction track concerns the prediction of pEC50 values for activation of human pregnane X receptor (PXR), measured using a cell-based reporter assay. PXR regulates the expression of drug-metabolizing enzymes and transporters, making its activation relevant to the assessment of drug interaction liabilities. The challenge combines primary activity measurements with auxiliary screening and PXR-null counter-assay data generated at Octant. Test compounds were selected as chemical analogues of potent screening hits, creating a prediction problem focused on structure–activity relationships within chemical series [10].

The activity track used 513 test compounds divided into Analog Set 1, containing 253 compounds, and Analog Set 2, containing 260 compounds. During Phase 1, participants received leaderboard feedback on Analog Set 1. Its labels were subsequently released for model refinement, while Analog Set 2 remained blinded until the challenge concluded. Final standings were determined by mean absolute error on Analog Set 2. The leading entry achieved an MAE of 0.406 and a Spearman correlation of 0.827, providing a reference for evaluation [18].

We replayed the original two-phase activity challenge within a 24-hour budget, preserving the phase-specific feedback and label-release sequence and evaluating the final submission on Analog Set 2. The five-agent run completed 19 submissions within the study cap of 46, fewer than the approximately 127 opportunities implied by the organizers’ announced schedule. In this retrospective replay, five-agent Codex achieved an MAE of 0.4049 on Analog Set 2 (Table 3), slightly outperforming the best reported challenge score of 0.4061 and placing the agentic result first by MAE relative to the 103 published final activity entries. Although these numerical comparisons do not establish a statistically significant difference from the leading challenge entry, the result extends the competitive performance observed on TDC and ExpansionRx to activity prediction within chemical series.

**Table 3.** Reference and agentic performance on the OpenADMET PXR Induction Challenge. Both rows evaluate activity prediction on Analog Set 2 (260 compounds). MAE determines the final activity ranking; Spearman correlation is a supplementary metric. Values in parentheses give the rank the configuration would occupy relative to the published final leaderboard, followed by the number of ranked entries (N).

| Endpoint | Metric | Best challenge | Tier1 min-max | CX-5 (rank/N) | Best agentic (rank/N) |
| --- | --- | --- | --- | --- | --- |
| PXR pEC50 Analog Set 2 | MAE ↓ | 0.4061 | 0.4061–0.4360 | 0.4049 (1/104) | 0.4049 (1/104) |
| PXR pEC50 Analog Set 2 | Spearman rho ↑ | 0.8269 | 0.7617–0.8269 | 0.8233 (3/104) | 0.8233 (3/104) |

Best challenge gives the best reported mean (matcha-croissant). Tier 1 min–max spans reported means across the organizers’ 28 Tier 1 entries (ranks 1–28); the Spearman range uses the same group, classified through statistical comparisons of MAE. These ranges are descriptive, not confidence intervals or thresholds for assigning new models to Tier 1. Only CX-5 results are reported for PXR. Bootstrap uncertainties are omitted from the table. Snapshot: 22 September 2026.
Configurations follow Table 1: CX, Codex with GPT-6 Astra (max). Numeric suffixes indicate agent counts.

**Table 4.**
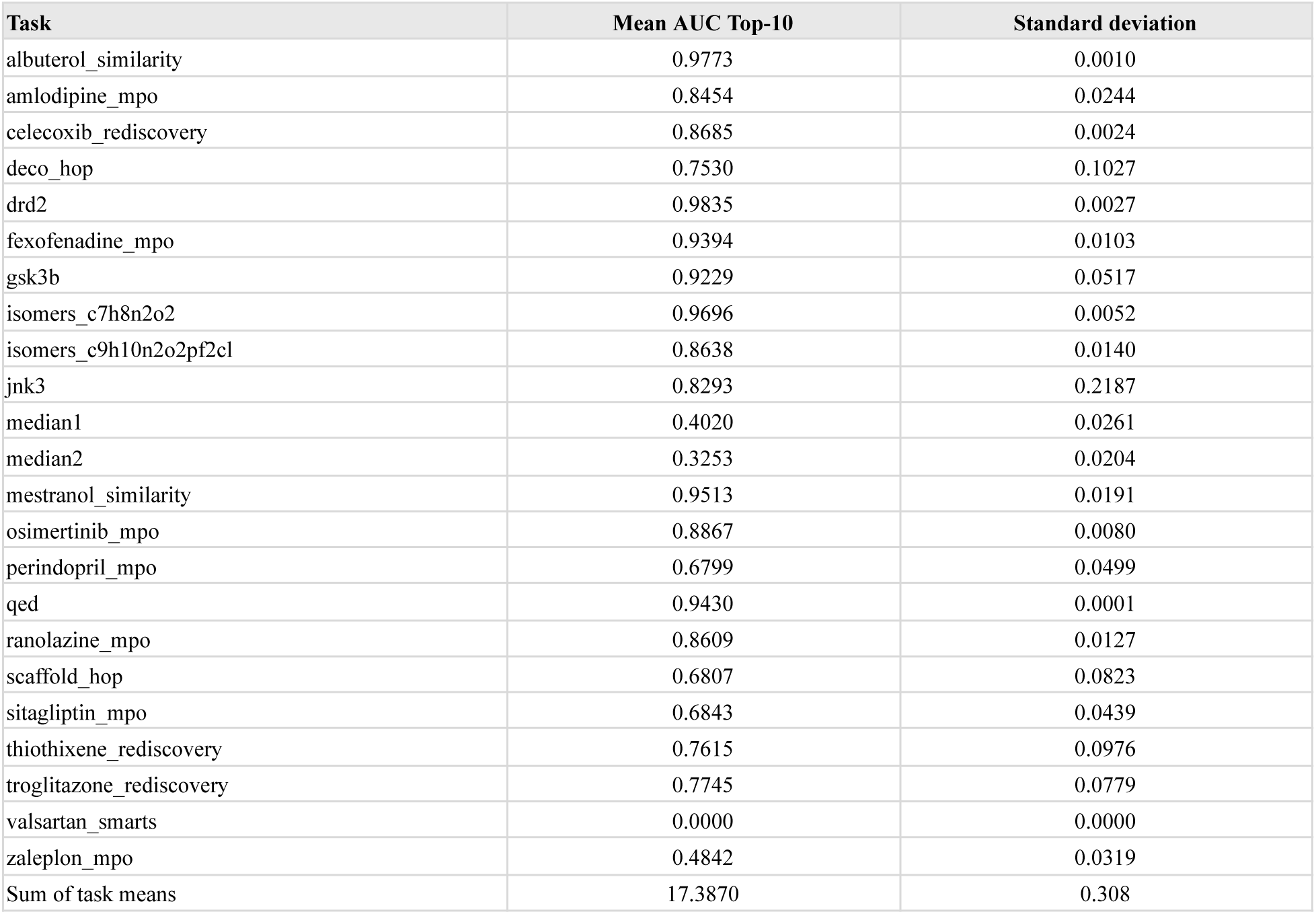
Results of the five-agent Codex running on GPT-6 Astra on the PMO benchmark. Values are the mean and standard deviation of the AUC Top-10 across the five seeds. The final row reports the sum of the 23 task means and the standard deviation of the per-seed sums.

The method developed by the agentic team used an ensemble that combined multilayer perceptrons (MLPs), directed message-passing networks, fine-tuned Uni-Mol, Uni-Mol2, various pre-trained molecular representation encoders [19–22], support-vector regressors, TabPFN [23, 24] and TabICL [25] regressors, and a LightGBM model [26], using additional residue-specific protein–ligand features from frozen Boltz-2 [27] and Protenix [28] models, followed by residual correction and partial refitting using the Analog Set 1 labels released to all participants at the original Phase 2 transition and made available at the corresponding point in our replay (detailed description in Supplementary Text S1.3).

### 2.4. Evaluation on the Practical Molecular Optimization benchmark

The Practical Molecular Optimization (PMO) benchmark [11] evaluates the ability to discover high-scoring molecules under a limited scoring budget. It comprises 23 tasks spanning molecular rediscovery, similarity, predicted biological activity, and multiproperty optimization. Performance is measured using AUC Top-10: the normalized area under the curve of the mean score of the ten best molecules found as a function of the number of oracle calls. This metric rewards both the quality of the discovered molecules and the efficiency with which they are identified. The overall score sums the task-level AUC values, with higher values indicating better performance.

We evaluated the same five-agent Codex configuration used for other benchmarks. Evaluation used the benchmark’s limit of 10,000 oracle calls per task and optimization run, with results averaged over five random seeds. Molecular pretraining was restricted to the supplied unlabeled ZINC250K collection [29]. During method development, the agents used feedback across zaleplon_mpo and perindopril_mpo tasks.

Five-agent Codex achieved an aggregate AUC Top-10 of 17.387 ± 0.308 under the original PMO protocol, exceeding the benchmark paper’s best result of 14.196, obtained by REINVENT [30], by 22.5%. Among the recent methods considered, Genetic GFN [31] provides the closest match to the original training-data and oracle-budget constraints and reports a lower aggregate score of 16.213. Other recent results were obtained under different experimental conditions: Mol-E [32], GenMol [33] and InVirtuoGen [34], including its variant without prescreening, use molecular pretraining data beyond ZINC250K, while RL-GFM [35] reports a 3,000-call budget rather than the original 10,000-call budget. Beyond differences in molecular training data, Mol-E incorporates pretraining supervision on properties that overlap with PMO objectives, including QED and components of several multi-property objectives. GenMol and the prescreened InVirtuoGen variant additionally use target-oracle evaluations before the measured optimization trajectory that are excluded from the nominal 10,000-call budget. Despite restricting molecular pretraining to the supplied unlabeled ZINC250K data, the agent-developed optimizer achieved performance comparable to the highest-scoring methods evaluated under the same 10,000-oracle-call protocol, with differences that are small relative to the reported run-to-run variability, while outperforming other recent methods under directly comparable conditions.

The agent-developed optimizer combined Gaussian-process-guided candidate selection, neural molecular generation, and genetic search, with molecular pretraining restricted to ZINC250K (see Supplementary Text S1.4 for details). Its performance illustrates the ability of the agentic workflow to integrate and refine established techniques into an effective molecular optimization method.

### 2.5. Cross-evaluation of agent-developed approaches

We evaluated the method developed by five-agent Codex for ExpansionRx on the 21 tasks of TDC ADMET, and the Codex-developed TDC approach on the nine ExpansionRx endpoints, in both cases applying the method as developed, without further modification. The ExpansionRx method transferred well, improving on the TDC+PR reference on 8 of 21 tasks. The TDC method transferred less successfully, obtaining an MA-RAE of 0.658, which would place 51st of 104 entries. Part of this gap may reflect the different development conditions: challenge participants could develop and iteratively refine their methods specifically for ExpansionRx using live half-test feedback, whereas we transferred a method developed for TDC with its modeling pipeline fixed, without allowing ExpansionRx-specific feedback-guided refinement.

Taken together, these cross-evaluation results indicate that agent-developed methods are not narrowly tuned to the benchmark they were developed on, although the degree of transfer varies with the direction and with how closely the two settings’ modeling demands align. Detailed results for both evaluations are provided in Supplementary Text S2.

## 3. Discussion

### 3.1. Novelty of agent-developed methods relative to leading solutions

To assess the novelty of the agent-developed methods, we compared them with existing leading solutions for each benchmark. The TDC method formed a distinct pipeline with custom molecular features, using a shared modeling approach across all endpoints. The ExpansionRx method showed substantial overlap with leading approaches, but the comparison did not establish reproduction or close adaptation of a particular top solution. For PXR, central modeling ideas had clear precedents, alongside a distinct implementation and a specific contribution in residue-specific protein–ligand feature design. For PMO, the agents combined prefix-guided SMILES generation, rank-weighted replay training, graph-based molecular edits, and Gaussian-process candidate selection, building on established components. Together, these findings support substantive work in modeling, feature engineering, and method integration, with the nature and strength of the novelty evidence varying across benchmarks.

### 3.2. Performance across evaluation settings

The TDC comparison tests whether agents can develop a reusable method across heterogeneous endpoints. Each endpoint’s reference comes from a separate, endpoint-specific method, so competitive performance under the single-method requirement suggests that autonomous method development can produce workflows with broad applicability. For this interpretation, the performance of individual agent runs is particularly informative: it measures what one complete method achieves, whereas selecting the best score across configurations measures the collective coverage of several development efforts. These comparisons address complementary questions about the reliability of a single solution and the benefit of exploring alternative agent configurations.

Notably, increasing agent count did not consistently improve performance across evaluation settings. On TDC, single-agent Claude Code outperformed its five- and ten-agent counterparts on 17 and 15 tasks, whereas on ExpansionRx five-agent Codex improved on the single-agent configuration on eight of nine endpoints. Team size should therefore be treated as a task-dependent design choice, with its benefit assessed under the available computational and time constraints.

PMO tests an additional capability: designing a procedure that determines which molecules to evaluate next. In the prediction benchmarks, performance depends primarily on extracting useful relationships from available measurements. In PMO, it also depends on allocating a limited scoring budget as new information becomes available. Competitive performance in this setting therefore provides evidence that agents can develop adaptive search procedures as well as predictive models. This distinction is relevant to research workflows in which learning and candidate selection are coupled, and the usefulness of a model depends partly on the subsequent evaluations it enables.

### 3.3. Limitations

The retrospective nature of these evaluations limits how the results should be interpreted. At the time of our experiments, the benchmark datasets and previously published approaches were publicly available. This is not unique to autonomous agents: human researchers working retrospectively on the same benchmarks may also be familiar with prior methods and results. For LLM agents, however, potential exposure through model pretraining cannot be directly determined or audited. The experiments therefore assess whether current agents can develop competitive methods under controlled benchmark conditions, rather than whether they can independently rediscover these approaches without any prior exposure. Prospective evaluations on previously unseen scientific problems will be important for assessing how these capabilities translate to real drug-discovery projects.

The retrospective challenge replays also differed in some respects from the original competitions. For ExpansionRx, the official training and full-test sets were retained, but the membership of the original half-test feedback set was unavailable, so feedback was provided on a fixed random half of the test set. The PXR replay preserved the original phase structure and evaluation population while compressing the competition timeline into a 24-hour run. The replay used study-specific compute limits and allowed up to 46 submissions, compared with approximately 127 submission opportunities under the original challenge schedule; these differences should be considered when making direct comparisons with challenge participants.

Finally, the reported statistical uncertainty primarily reflects variability in evaluation across test compounds rather than variability in autonomous method development. Independent agent-development runs were not repeated sufficiently to estimate how consistently the systems would arrive at the reported solutions. In the PXR experiment, three of the five agents also terminated because of model-service capacity errors, resulting in unequal participation across agents. Repeated independent runs would be needed to characterize the robustness and run-to-run variability of autonomous research performance.

## 4. Conclusion

Across four molecular modeling settings, LLM agents autonomously developed methods that surpassed leading human-developed methods on several benchmarks and remained highly competitive on others. The results span heterogeneous ADMET prediction, temporal generalization during lead optimization, activity prediction within chemical series, and molecular optimization under a limited oracle-call budget. Together, they show that current agents can carry out substantial parts of molecular model development, including representation selection, model training, feature engineering, ensembling, calibration, and adaptive molecular search.

At the same time, retrospective benchmarks should be viewed as controlled tests of capability rather than as the ultimate measure of scientific usefulness. The more consequential test is therefore prospective: whether autonomous agents can tackle previously unseen scientific problems, generate useful predictions and methods without retrospective exposure, and ultimately produce results that hold up in new experimental measurements. Prospective studies that connect autonomous computational research to experimental validation will be essential for establishing how far these systems can contribute to real-world drug discovery.

## 5. Methods

### 5.1. Agent task and evaluation protocol

For each benchmark, agents were evaluated under the official protocol: the prescribed data splits, the prescribed metrics, and the published scoring implementation where one was available.

The benchmark execution framework creates a fresh temporary user account with restricted permissions for each run. All agents within a run share this account and can access their workspaces, teammates’ research files and the shared research log. The account has no administrative privileges, and agent processes are prevented from acquiring additional privileges. Evaluation services operate separately under the experiment operator’s account. Private evaluation data and the records used to enforce submission limits are stored with permissions that deny access to the agent account. When the run ends, the system stops its remaining processes, removes the temporary account and returns ownership of the research outputs to the operator. For each run, the agentic team shared a single allocation of one NVIDIA H100 GPU, 16 virtual CPUs, and 200 GiB of memory.

Humans provided no scientific guidance, code modifications, model selection, or other intervention. Human involvement was limited to setting up the evaluation environment, preparing the task prompt, and auditing the outputs and agentic traces after completion.

The LLM prompts used for each benchmark are provided in Supplementary Text S3.

#### TDC ADMET Benchmark Group

We evaluated 21 of the 22 datasets in the TDC ADMET Benchmark Group, covering absorption, distribution, metabolism, excretion, and toxicity. Bioavailability_Ma was excluded because of unresolved inconsistencies in its TDC leaderboard scores [36], which complicated comparison with previously reported methods. The tasks involve predicting molecular properties from chemical structures provided as SMILES strings and include both regression and binary classification.

The benchmark uses predefined scaffold-based partitions, with approximately 20% of each dataset reserved for a fixed test set. This splitting strategy groups molecules by their core structures to assess generalization across chemical scaffolds. Depending on the endpoint, regression performance is evaluated using mean absolute error or Spearman’s rank correlation, while classification performance is evaluated using the area under the receiver operating characteristic or precision–recall curve. The official leaderboard protocol ranks the methods based on the mean score across at least five independent runs evaluated on the same test set.

We tasked LLM agents with developing predictive methods for 21 TDC ADMET benchmarks, aiming to improve on published results. Within each run, agents were required to develop a single method applicable across all 21 tasks, whereas each task’s reference score was taken from the best-performing previously reported method for that endpoint. Each agent run received a 15-hour wall-clock budget covering research, implementation, training, validation, evaluation, and report preparation. Agents were allowed three scoring submissions, each of which could evaluate a subset or all benchmarks. Performance was assessed separately using each benchmark’s prescribed metric, without an aggregate score.

Agents were instructed to use the supplied training–validation partitions for model development and selection and to document their validation procedure. The instructions provided TDC’s default scaffold validation procedure, targeting an 87.5%/12.5% training–validation split. Evaluation used the unchanged official scaffold test partitions, and the scorer verified that the supplied data files had not been modified. Although test labels were present in the distributed files, agents were explicitly prohibited from using test labels to guide preprocessing, training, hyperparameter tuning, or model selection. Agent execution traces were reviewed for violations of these restrictions. Results intended for leaderboard comparison required at least five independently trained runs, with the mean and standard deviation reported for each benchmark.

Agents were required to provide saved models and preprocessing artifacts, a standardized prediction interface, and sufficient code and dependency information to reproduce the results. A technical report documented the method, its contribution relative to prior work, experimental procedures, results, unsuccessful experiments, and limitations. Comparisons used a dated leaderboard snapshot, with relevant differences in evaluation conditions disclosed. Agents were also required to identify external datasets and pretrained models and report any known or possible exposure to TDC test molecules or labels.

#### OpenADMET ExpansionRx Challenge

In a retrospective replay of the ExpansionRx challenge, we evaluated Codex (GPT-6 Astra, maximum reasoning effort) in single-agent and five-agent configurations with the same data, feedback protocol and resource limits. The replay retained the official temporal split: 5,326 earlier-stage training molecules with 25,265 observed endpoint measurements, and 2,282 later-stage test molecules supplied only as identifiers and SMILES strings. The raw training data were also available, and missing endpoint measurements were kept as missing values. Each configuration had a 24-hour wall-clock budget (covering research, training, local validation and report preparation), at most 30 scoring submissions. These budgets were defined for this study rather than taken from the original challenge, in which human participants were limited to one submission per day over a 12-week window (27 October 2025 to 19 January 2026), i.e., at most about 85 submissions per account.

The replay followed the organizers’ feedback protocol (live half-test feedback plus one intermediate and one final full-test evaluation) on a compressed clock. Each accepted submission covered all 2,282 test compounds and all nine endpoints. Half-test feedback reported the official metrics on a fixed private subset of 1,141 test molecules, sampled randomly once at setup because the original subset’s membership was unavailable. Agents were free to choose their local validation strategy and could use the permitted aggregate feedback for model selection and calibration. Following the organizers’ intermediate full-test evaluation, the replay provided one frozen full-test score for the latest accepted submission at 3/7 of the 24-hour budget, approximately 10 h 17 min after registration. At the deadline, all worker processes were stopped and the latest accepted submission was evaluated on the full test set, even if an earlier submission had received better half-test feedback. The five-agent run lasted 18 h 09 min 15 s, with all 30 submissions accepted before the deadline; submission 20 received the intermediate full-test evaluation and submission 30 was the final entry. The required predictions and methodology reports were archived.

Scoring used the official evaluation code. LogD was scored on its original scale; the other eight endpoints were scored after transforming observations and predictions as *log_10_* (*max*(*y*, 0) + 1), while submitted CSV files remained in the original units. Predictions were matched to labels by molecule identifier and scored only where a label was observed. An endpoint’s relative absolute error (RAE) is its MAE divided by the mean absolute deviation of the observed labels from their mean, and the overall score (MA-RAE) is the unweighted mean of the nine endpoint RAEs. Uncertainty was estimated from 1,000 bootstrap resamples (NumPy seed 0), with the RAE denominators and MA-RAE recomputed within each resample; we report the bootstrap mean ± sample standard deviation and, consistent with the official evaluator, use the bootstrap mean for leaderboard comparisons. An independent recalculation reproduced the five-agent full-test MA-RAE of 0.537814 ± 0.006792 (point estimate on the non-resampled test set: 0.537197).

Reference scores were taken from the official full-test leaderboards (the per-endpoint leaderboards and the FINAL tab; 22 September 2026 snapshot, Table 2). For each endpoint, the reference is the lowest published mean MAE; the overall reference is the lowest published MA-RAE achieved by a single complete submission, rather than a combination of the best per-endpoint results. The two configurations were ranked separately by retrospectively inserting each score into the published ranking of 103 eligible entries; we do not assign either to the organizers’ statistical groups or claim statistically significant differences from neighboring entries.

#### OpenADMET PXR Induction Challenge

We evaluated five-agent Codex on the activity track in a retrospective replay of the official challenge. The target was primary human PXR induction pEC50, predicted directly in the supplied units. The replay retained the original training resources, all 513 blinded activity compounds, and their partition into Analog Set 1 (253 compounds) and Analog Set 2 (260 compounds). Initial inputs included 4,139 primary dose-response measurements, the counter-assay and single-concentration training tables, and optional structure-track specifications. We used the official data available during Phase 1, including the organizers’ corrections, and released additional measurements at the phase transition.

Five Codex workers using GPT-6 Astra at maximum reasoning effort shared a maximum of 24 hours and 46 submissions across both phases. These were study-specific budgets. Under the organizers’ announced schedule of one submission every four hours during the first week and one per day thereafter, human participants would have had approximately 127 submission opportunities per track over the full challenge. The challenge’s phase sequence was retained, with its calendar compressed into the local runtime: the handover occurred after 14 h 41 min 05 s, preserving the corrected May 26 Pacific cutoff as a fraction of the April 1–July 1 prediction period. Upload cooldowns were also scaled to the local clock.

During Phase 1, each submission contained predictions for all 513 compounds, but live feedback covered only Analog Set 1, with RAE as the primary live metric. At handover, the latest accepted Phase 1 submission received one frozen aggregate evaluation on all 513 compounds, and Set 1 labels plus the original crude and semi-pure HTChem measurements were released. Phase 2 permitted their use for training and validation, while new submissions returned acceptance receipts without scores. Thus, Set 1 was development data, and Set 2 contributed the single authorized aggregate checkpoint but no released labels or subsequent live feedback. At closure, the latest accepted submission with a timely methodology report was evaluated on Set 2 alone, ranked by MAE. These transitions follow the organizers’ published procedure.

Scoring followed the original evaluator, using 1,000 bootstrap resamples with NumPy seed 0 and reporting the mean and sample standard deviation across resamples. No additional logarithmic transform or retrospective weighted-MAE proposal was applied. Independent recalculation reproduced the final MAE of 0.40488 ± 0.02923 and Spearman correlation of 0.82328 ± 0.02812. The ordinary, unbootstrapped MAE was 0.40471; leaderboard comparisons use the bootstrap mean.

The agent’s PXR run made 19 submissions, comprising 17 in Phase 1 and two in Phase 2, and closed after 20 h 40 min 49 s. The final predictions and methodology report were accepted before closure. The ranking is a retrospective comparison with the published final leaderboard.

#### Replay integrity and verification

Retrospective challenge replays used private evaluation services that withheld test labels and enforced the specified submission and feedback protocols. Because challenge labels and participant solutions were publicly available at the time of the experiments, agents were explicitly prohibited from retrieving post-challenge labels, final rankings, or participant implementations. Independent review of retained agent interaction records found no direct acquisition or use of hidden evaluation labels and no evidence of copying from the examined participant solutions.

Final ExpansionRx and PXR predictions were independently reconstructed from archived model outputs and rescored using the original evaluation implementations, reproducing the reported metrics. These audits cannot exclude exposure through the pretrained LLMs or pretrained molecular models.

#### Practical Molecular Optimization benchmark

We evaluated a team of five Codex agents (GPT-6 Astra, maximum reasoning effort) on the Practical Molecular Optimization benchmark. The agents received an evaluation interface and the ordered ZINC250K pool, containing 249,455 unlabeled SMILES strings. Molecular pretraining and initialization assets were restricted to this pool. Consulting published methods was permitted. Research had no fixed deadline. The run took approximately 24.5 hours to complete.

Model and hyperparameter selection for the reported optimizer used the two designated tuning tasks, zaleplon_mpo and perindopril_mpo. Final evaluation covered all 23 PMO objectives using the selected learning and search settings, with each trial starting from the same unconditional molecular generator pretrained on ZINC250K.

Final evaluation comprised all 23 objectives and five optimization seeds (0, 1, 2, 3 and 5), giving 115 trials with at most 10,000 counted oracle evaluations each. All target scores, including those used for initialization and online surrogate training, passed through the supplied scoring service. A valid molecule consumed one call on its first occurrence after canonicalization; subsequent requests for the same canonical molecule returned its cached score without another charge. Invalid SMILES returned zero without consuming a call. Each trial began with a fresh oracle cache and the same permitted starting assets, and could learn from its own counted observations. The reported result came from one complete evaluation with a frozen method, configuration and dependency specification; results were not assembled from different method versions or selected seeds.

Evaluation used the local corrected protocol based on the original PMO implementation and a pinned scoring environment with PyTDC 0.3.6 [37] and RDKit 2022.3.5 [38]. The sitagliptin_mpo scoring function was corrected using TDC commit d566b108 [39]. No additional synthetic-accessibility, diversity or novelty criterion was added to the primary score.

The primary metric was normalized AUC Top-10. For each trial, the evaluator calculated the mean of the ten highest scores observed so far at 100-call intervals, beginning at zero and including the final partial interval. It integrated this curve using the trapezoidal rule and divided the area by 10,000. Successful early termination extended the final achieved Top-10 mean through the remaining budget; failed or interrupted trials could not contribute to a complete benchmark aggregate. Scores were averaged across the five seeds within each objective and then summed without weighting across the 23 objectives, yielding an overall score between 0 and 23. Aggregate variability was calculated from the five totals obtained by summing all task scores at each seed.

## Acknowledgments

We thank Natalie Ma and W. Patrick Walters for their careful reading of the manuscript and for their constructive comments and suggestions.

## Supplementary Text

## S1. Methods developed by agents

This section describes the top-performing methods developed by the agents.

### S1.1. Five-agent Codex solution for TDC ADMET benchmarks

All endpoint predictors are TabM networks: multilayer perceptrons that share weight matrices across k internal members, each member having its own modulation and output parameters. A configuration fixes three things: (i) the input representation, (ii) the network settings (width, depth, k, dropout, learning rate), and (iii) the training objective. The prediction for each endpoint is a combination of the outputs of one or more configurations. Combinations were selected by scaffold-split validation on the training data.

For the network and training, the default settings are width 384, depth 3, k = 8, dropout 0.2, and learning rate 5 × 10^-4^ with AdamW. Configurations with descriptor-only features instead use width 256, k = 16, dropout 0.1, and learning rate 10^-3^. The default objectives by metric are:

- binary cross-entropy for classification endpoints;
- Huber loss (β = 0.5) on standardized targets for MAE endpoints;
- mean squared error on standardized log_10_ targets for Spearman endpoints.

Each configuration used an input representation from the list provided in Table S1. Inputs are pre-computed molecular descriptors, frozen pretrained encoders (their embeddings are used as fixed inputs and not fine-tuned), or both.

**Table S1.**
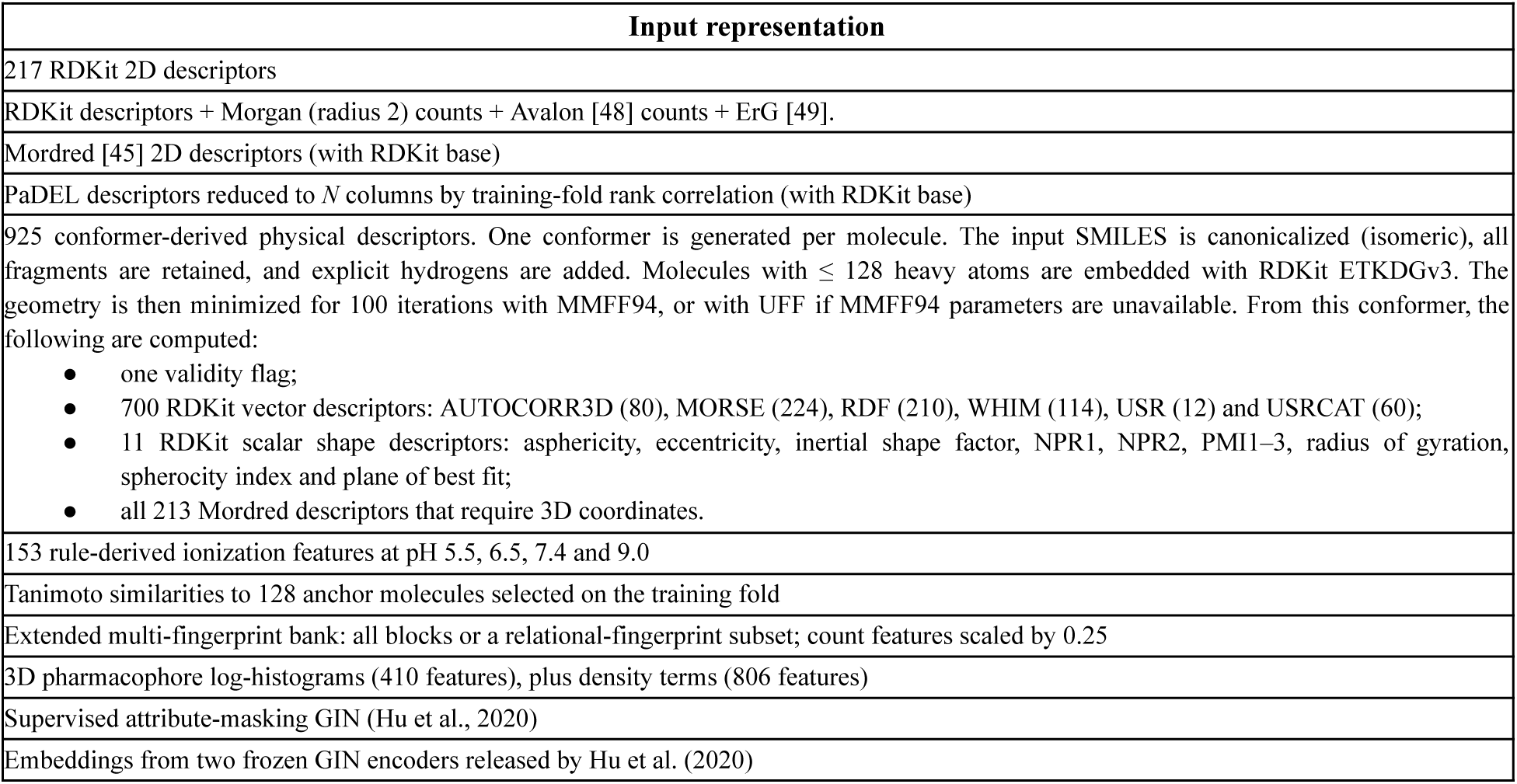
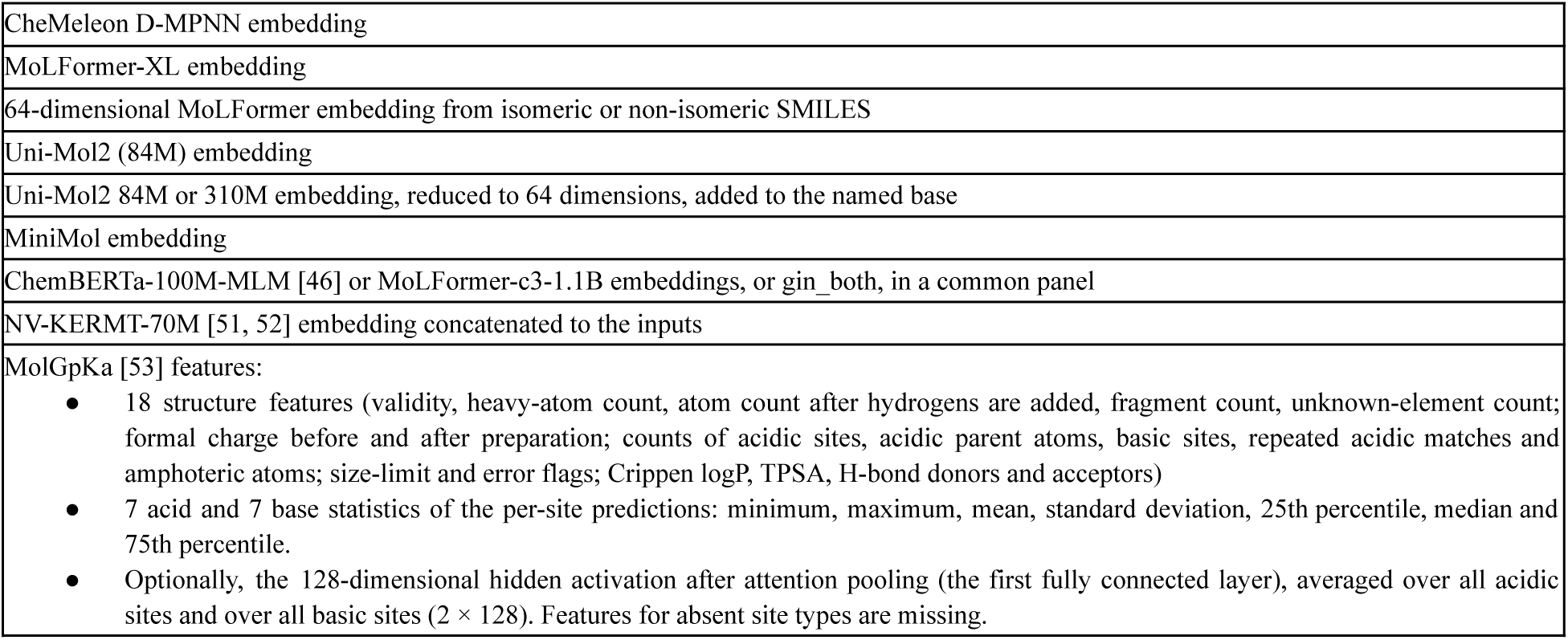
Input representations used in agent-developed TabM ensembles for TDC.

### S1.2. Five-agent Codex solution for the ExpansionRx challenge

The predictor combines TabM networks, multilayer perceptrons (MLPs), directed message-passing networks, fine-tuned MiniMol and Uni-Mol models, and molecular kernel regressors. Each of the nine ADMET regression endpoints uses its own ensemble (Table S2). Configurations differ in molecular inputs, model family, network dimensions where applicable, and training objective. Tabular models use descriptors, fingerprints, frozen pretrained embeddings and public-assay predictions (Table S3); graph and 3D models additionally learn from molecular structure.

TabM configurations use three 512-dimensional layers, k = 8 or 16 internal members, dropout 0.15, and AdamW with learning rates of 6 × 10^-4^ −10^-3^. MLPs use three hidden layers of width 512 or 1,024. Directed message-passing networks use hidden width 500 and message depth 5. The original Uni-Mol encoder is fine-tuned, with predictions averaged over five conformers at inference.

Training uses 5,326 molecules with partially observed labels. LogD remains untransformed; the other endpoints use log_10_(max(y, 0) + 1). Targets for the neural network models are subsequently standardized within each fitting partition. TabM, directed graph and Uni-Mol models use masked, task-weighted Smooth-L1 losses [40]; MLPs use a masked, task-weighted smooth absolute-error loss,√(*e*^2^ + 0.1^2^) − 0.1, where e is the standardized prediction error. Kernel models predict ADMET properties from molecular similarity using ridge or support-vector regression, with selected variants accounting for censored measurements or sharing information across endpoints.

Selected models used the relationship between the MBPB and MGMB binding endpoints to supplement missing training labels or link their predictions. For TabM, a linear MBPB-to-MGMB mapping is fitted by Huber regression to paired, log-transformed measurements within each fitting partition. This mapping estimates missing MGMB training labels from observed MBPB, while its inverse estimates missing MBPB labels from observed MGMB. These pseudo-labels receive half the loss weight of measured labels; observed and validation labels remain unchanged. In graph models, the MGMB output combines 20% of its direct prediction with 80% of an estimate derived from predicted MBPB using a separately fitted linear mapping, with both terms expressed in standardized MGMB coordinates. For efflux prediction, an auxiliary reverse-permeability target is calculated as forward permeability × efflux wherever both measurements are available. Models trained on forward and reverse permeability estimate efflux as predicted reverse permeability divided by predicted forward permeability, after reversing the target transformations.

Model selection uses five-fold row or parent-grouped cross-validation, ordered holdouts and structure-based holdouts. Neural checkpoints are selected separately for each endpoint, and predictions from fold-trained models and full-data refits are combined on the log-transformed scale and back-transformed afterwards. The final ensemble includes endpoint-specific weights, out-of-fold residual corrections, fixed offsets, and a variance adjustment that rescales each prediction about the mean over the molecules predicted together in one batch. Predictions are constrained to nonnegative values in original units for all endpoints except LogD, with binding percentages additionally capped at 100%. Final model and calibration choices also use repeated aggregate feedback from a 1,141-molecule validation subset included in the 2,282-molecule full-test evaluation.

**Table S2.**
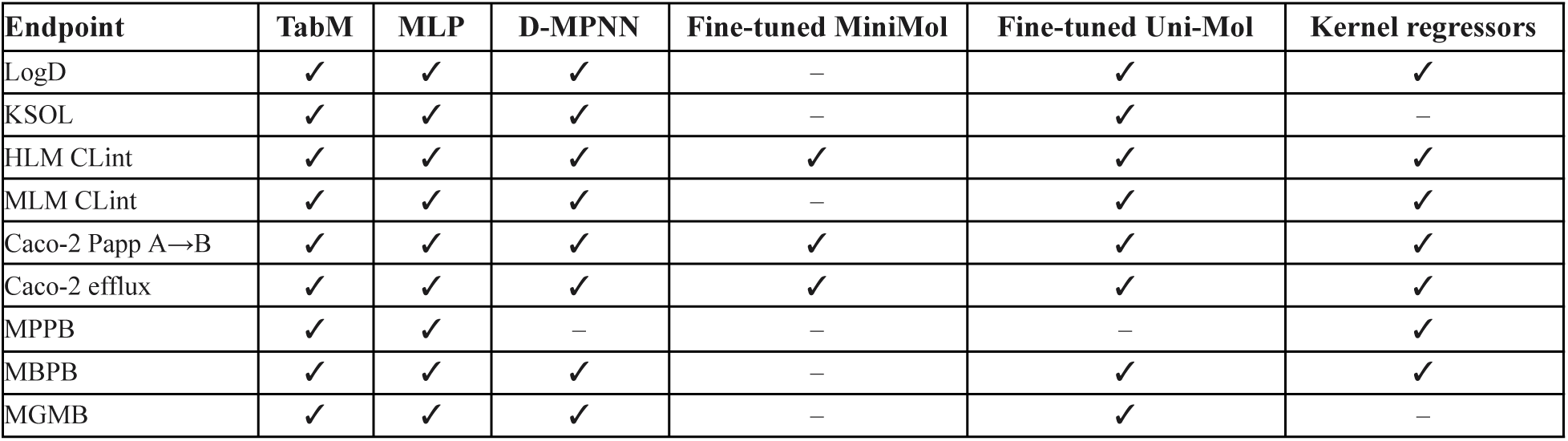
Model families contributing to each endpoint ensemble. A checkmark indicates that the model family contributes endpoint predictions. MLP configurations may use frozen pretrained embeddings and public-assay predictions; the MiniMol and Uni-Mol columns denote separately fine-tuned predictors. KSOL additionally uses source-assay and spline residual corrections. HLM, Papp and efflux use source-assay, spline and kernel residual corrections. Efflux also incorporates predictions derived from the forward/reverse-permeability kernel models.

**Table S3.**
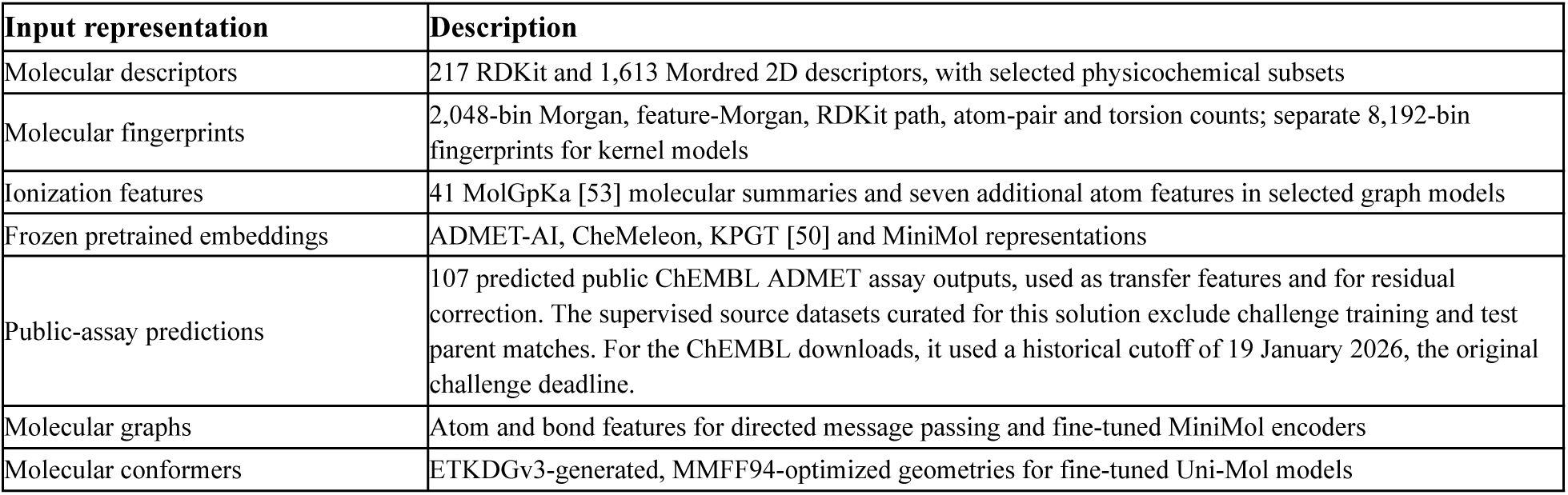
Molecular inputs used across the ensemble.

| Input representation | Description |
| --- | --- |
| Molecular descriptors | 217 RDKit and 1,613 Mordred 2D descriptors, with selected physicochemical subsets |
| Molecular fingerprints | 2,048-bin Morgan, feature-Morgan, RDKit path, atom-pair and torsion counts; separate 8,192-bin fingerprints for kernel models |
| Ionization features | 41 MolGpKa [53] molecular summaries and seven additional atom features in selected graph models |
| Frozen pretrained embeddings | ADMET-AI, CheMeleon, KPGT [50] and MiniMol representations |
| Public-assay predictions | 107 predicted public ChEMBL ADMET assay outputs, used as transfer features and for residual correction. The supervised source datasets curated for this solution exclude challenge training and test parent matches. For the ChEMBL downloads, it used a historical cutoff of 19 January 2026, the original challenge deadline. |
| Molecular graphs | Atom and bond features for directed message passing and fine-tuned MiniMol encoders |
| Molecular conformers | ETKDGv3-generated, MMFF94-optimized geometries for fine-tuned Uni-Mol models |

### S1.3. Five-agent Codex solution for PXR Induction challenge

The method predicts PXR agonist potency as *pEC*_50_ =− *log*_10_(*EC*_50_/*M*) using an ensemble of 33 components. These comprise multilayer perceptrons (MLPs), directed message-passing networks, fine-tuned Uni-Mol, Uni-Mol2, CheMeleon and MoLFormer encoders, support-vector regressors, TabPFN and TabICL regressors, and a LightGBM model. Initial training uses 4,139 primary-assay molecules, supplemented with counter-assay and single-concentration measurements as auxiliary targets. In Phase 2, the released labels for Analog Set 1 (253 compounds) are used to retrain selected ensemble components and fit a correction for the ensemble’s prediction errors. Final performance is evaluated by MAE on Analog Set 2 (260 compounds). The input representations used in this solution’s models are summarized in Table S4.

**Table S4.** Input representations used in five-agent Codex solution for PXR Induction challenge.

| Input representation | Role |
| --- | --- |
| 7,335-dimensional neural fingerprint vector: 2,048-bin radius-2 and radius-3 Morgan counts; 1,024-bin feature-Morgan, atom-pair and torsion counts; 167 MACCS bits. Counts are clipped at 255 and transformed with $\log(1+x)$. | Dense neural inputs |
| 217 ordered RDKit descriptors; additional Mordred 2D descriptors in selected kernel components | Neural, graph, tree, tabular and kernel inputs with component-specific preprocessing |
| 4,096-bin radius-2 Morgan counts; separate 8,192-bin fingerprint representations for kernel models | SVD-reduced tabular features, tree inputs and molecular similarity kernels |
| Frozen MoLFormer pooled embeddings (768), ChemFM mean-token embeddings (3,072), Uni-Mol2 CLS embeddings (768), CLAMP hidden embeddings (2,048) and CheMeleon atom-pooled embeddings (2,048) | Transfer representations for MLPs and tabular foundation regressors |
| Molecular graphs with 133 atom features and 14 directed-bond features | Directed message passing; pretrained graph encoders use their own feature definitions |
| Seed-42 RDKit heavy-atom conformers with MMFF optimization when successful | Frozen and fine-tuned Uni-Mol representations; ligand inputs to protein-conditioned extraction |
| Boltz core (1,280), pocket embedding summary (768), pocket residue structural embedding (3,968) and whole-protein structural embedding (37,504); Protenix pocket residue structural embedding (3,968) | Per-residue structural embeddings are used in the neural networks, RBF kernel similarities in the support-vector regressors, and 64-component PCA reductions in the tabular models. Molecules with no computed structure have that block zero-filled and carry an explicit availability flag. |

The principal MLP architecture projects concatenated fingerprints and pretrained embeddings to 512 units and descriptors to 128 units. Their concatenation passes through two additional 512-dimensional layers and a linear multitask output layer. Hidden layers use LayerNorm, SiLU and dropout 0.25. Training uses AdamW optimizer with learning rate ^−4^, weight decay 0.01, batch size 256, cosine learning-rate decay and up to 160 epochs. Primary potency uses absolute or squared error, depending on the component; auxiliary targets use squared error. The primary loss weight is 1, with total weights 0.15, 0.1 and 0.75 assigned to primary-assay auxiliary endpoints, counter-assay endpoints and single-concentration screening endpoints, respectively; the screening endpoints are kept separate by experiment and concentration. Losses are masked for missing observations, and targets for the neural networks are standardized within each fitting partition. Multitask MLP epochs balance retained primary rows with an equally sized sample of auxiliary-only rows. Predictions average the five epochs with lowest primary validation MAE and, where specified, multiple seeds.

The graph ensemble combines a width-300, depth-3 directed message-passing network, width-512, depth-5 networks, and a teacher-augmented width-300 network, with internal weights 0.2, 0.6 and 0.2. Mean-pooled atom representations are concatenated with RDKit descriptors. These models predict primary potency, assay efficacy, counter-assay endpoints and three concentration-specific screening responses using a masked, task-weighted Smooth-L1 loss with parameter 0.5. Training uses AdamW optimizer with learning rate 10^−3^, weight decay 10^−5^, batch size 64 and a 100-epoch schedule. The teacher branch additionally uses screening-derived pseudo-potencies with observation weight 0.15. Uni-Mol2 components fine-tune the 84-million-parameter, 12-layer, 768-dimensional encoder for primary-only or eight-task prediction. Encoder learning rates are 3×10^−5^5×10^−5^, with tenfold higher head learning rates, batch sizes 32–64, and 25–35-epoch horizons. Their selected inference states use an exponential moving average capped at decay 0.99.

Protein-conditioned features are extracted from frozen Boltz-2 and Protenix model trunks, using the June 2025 checkpoint for Protenix. Inputs use the 293-residue PXR construct and a 1NRL chain-A template [41]; an additional Boltz branch uses the 1ILH template [42]. The 31-residue pocket is defined by proximity within 6 Å of the native 1NRL ligand. Averaging the forward protein-to-ligand pair representation over ligand tokens yields 128 values per residue, giving a 3,968-dimensional Boltz pocket residue embedding. Additional components use pooled single/pair features or per-residue structural embeddings for all 293 residues. Boltz extraction uses its initial trunk without additional recycling; Protenix uses one trunk cycle. Diffusion and affinity heads are not run. These frozen representations are supplied to supervised MLPs, kernel models and tabular regressors.

Kernel components combine fingerprint Tanimoto similarities with radial-basis kernels over normalized RDKit, Mordred or pocket residue-embedding features. Support-vector regression uses ɛ = 0. 1 and selected *C* values of 1, 3 or 10. TabPFN v2/v2.5 and TabICL v2 [47] use descriptors, fold-fitted PCA reductions of frozen molecular embeddings and per-residue structural embeddings, and SVD reductions of log-count fingerprints. Selected PCA widths are 48, 64 or 128 per block. TabPFN uses 4 or 16 internal estimators and TabICL uses 8; conditional medians are used except for one TabPFN v2 mean-output component. Their pretrained parameters remain fixed while retained labeled molecules form the prediction context.

Original model selection uses five-fold, target-stratified cross-validation grouped by canonical achiral molecular identity. Within each fold, primary targets, auxiliary supervision, scalers and dimensionality reductions all see only retained rows. Most components predict by averaging the five fold fits; the tree and molecular-fingerprint SVR bundle refit on all data. Component weights come from nonnegative MAE optimization on out-of-fold predictions, averaging leave-one-fold-out fits under global and potency-neighbor-balanced objectives. The resulting original predictor b(x) is a weighted average of the 33 component predictions, with nonnegative weights summing to one, up to recorded floating-point precision.

Analog Set 1 molecules are assigned to their nearest original-training molecule using binary radius-2 Morgan Tanimoto similarity. This produces 49 parent groups, partitioned into five folds. Each validation fit retains all original labels and only the other four released folds. Fifteen component slots are updated, corresponding to 14 distinct predictors because two Boltz slots share one updated model. MLP, graph and Uni-Mol2 refits each average three fixed seeds, with snapshot schedules derived from original development fits rather than selected on the held-out released fold.

A median residual regressor is fitted to *y* − *b*(*x*) using the frozen prediction *b*(*x*) and four family contrasts *F_k_* (*x*) − *b*(*x*), where each *F_k_* is the within-family weighted mean for molecular MLPs, protein-conditioned MLPs, supervised 3D models or tabular foundation models. Features are standardized on retained released rows. Calibration uses quantile 0.5, an intercept and an *L_1_* coefficient penalty of 0.05. With residual prediction *g*(*x*), ∼ updated component outputs *f_j_* (*x*), and updated-slot set *J*, the final predictor is

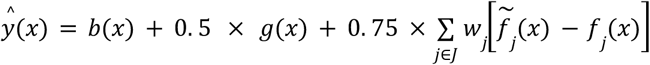

Original weights and calibration features remain fixed when applying the updates. The update strength 0.75 is selected from {0, 0. 1, 0. 25, 0. 5, 0. 75, 1} as the lowest-MAE candidate that improves all five released-fold MAEs relative to the calibrated frozen baseline. Final updated models use all 4,392 primary labels; final calibration uses all 253 released residuals.

Evaluation uses the 260 remaining molecules, ordered by molecule identifier. The reported MAE is the mean over 1,000 bootstrap samples of 260 rows drawn with replacement using NumPy’s default random generator with seed 0.

### S1.4. Five-agent Codex solution for PMO benchmark

The method combines a recurrent SMILES generator, graph-based molecular edits, and a Gaussian-process (GP) surrogate. The generator is a three-layer gated recurrent unit network [43] with 384 hidden units per layer and 128-dimensional token embeddings; overall parameter count is 2,393,397. It is pretrained for 20 epochs using next-token cross-entropy on a corpus derived from unlabeled ZINC250K molecules, each represented by one canonical and two randomized SMILES. Pretraining uses Adam with batch size 512 and initial learning rate 10^-3^, multiplied by 0.92 after each epoch with a floor of 10^-4^.

Each optimization trial starts from the pretrained checkpoint and an initial sample of 100 ZINC entries, filtered for validity and duplicates. At each iteration, the generator receives four Adam optimizer updates with batch size 64 and learning rate 2 × 10^-4^. Training examples are sampled from the 100 highest-scoring molecules with weights inversely proportional to their zero-based rank plus 10. Each replay example uses randomized SMILES with probability 0.75 and canonical SMILES otherwise. The training objective is L = E_s_[(NLL_θ_(s) − NLL_prior_(s) + 60 × R(s))^2^], where R(s) is the observed reward and each NLL is the sum of non-padding token negative log-likelihoods, including the end-of-sequence token [30]. The pretrained prior remains frozen.

Each iteration generates 96 neural proposals and makes 48 graph-edit attempts using crossover and mutation. Neural proposals use a prefix from a randomized replay-parent SMILES with probability 0.75. The retained prefix fraction is drawn uniformly from 0.35–0.90; the number of retained tokens is the fraction multiplied by the parent’s token count, rounded down. Invalid, duplicate, and previously evaluated candidates are removed. The graph edits use the Graph GA reaction-template operators [44].

The GP is fitted to at most 1,000 observed molecules. When the history exceeds this limit, each fit uses the 333 highest-scoring molecules and 667 molecules sampled without replacement from the remainder. Rewards are centered by their mean, and diagonal regularization of 0.001 is added to the kernel matrix. The kernel combines fingerprint Tanimoto similarity and a radial-basis kernel over physicochemical descriptors:

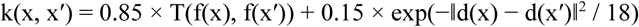

where f is the Morgan count fingerprint and d is the scaled descriptor vector in Table S5. The fingerprint kernel uses the dot-product form T(f, g) = (f · g) / (ǁfǁ^2^ + ǁgǁ^2^ − f · g).

Up to 24 candidates are selected greedily using an upper confidence bound, μ(x) + 0.05 × σ(x), where σ(x) is the GP posterior standard deviation. After selecting a candidate x*, the acquisition value of each remaining candidate x is reduced by 0.015 × k(x, x*)^6^. This similarity penalty accumulates across selections to encourage diversity within the batch.

The configuration and pretrained checkpoint were selected using the development objectives **zaleplon_mpo** and **perindopril_mpo** and then fixed across all 23 PMO objectives. Each objective uses five seeds (0, 1, 2, 3, and 5) and a budget of up to 10,000 distinct valid oracle evaluations. The top-10 reward curve is integrated using the trapezoidal rule at 100-call intervals, including the final observed endpoint. Successful early stops are extended horizontally to the 10,000-call horizon, and the area is normalized by 10,000. Performance is the sum across objectives of the five-seed mean normalized area under this curve.

Table S5 describes the input representations used in this solution’s models.

**Table S5.** Input representations used in five-agent Codex solution for PMO.

| Input representation | Role |
| --- | --- |
| Tokenized SMILES with a 53-token vocabulary | Pretraining, replay training, and generation |
| RDKit molecular graphs with stereochemistry removed | Crossover, mutation, and validity checks |
| 2,048-dimensional Morgan count fingerprints, radius 2 | Structural component of the GP kernel |
| 18 scaled physicochemical descriptors, in order: atom counts for C, N, O, F, P, S, Cl, Br and I; attached-hydrogen count; aromatic-atom count; ring count; rotatable-bond count; hydrogen-bond donor count; hydrogen-bond acceptor count; TPSA; Crippen logP; and exact molecular weight. The first 11 entries are multiplied by 0.25, TPSA is divided by 20, and molecular weight is divided by 40; the remaining entries are unscaled. | Property component of the GP kernel |

## S2. Cross-evaluation performance of agent-developed approaches

### S2.1. TDC ADMET performance of the five-agent Codex solution developed for ExpansionRx

Table S6 reports the performance of the agent-developed ExpansionRx method on the TDC ADMET Benchmark Group. We retained the original modeling pipeline without modification. Because ExpansionRx includes only regression tasks, TDC classification tasks were modeled as regression on binary (0/1) labels, preserving the original regression models and loss functions. Their continuous predictions were evaluated directly using ROC-AUC or average precision, according to each task’s prescribed metric.

**Table S6.**
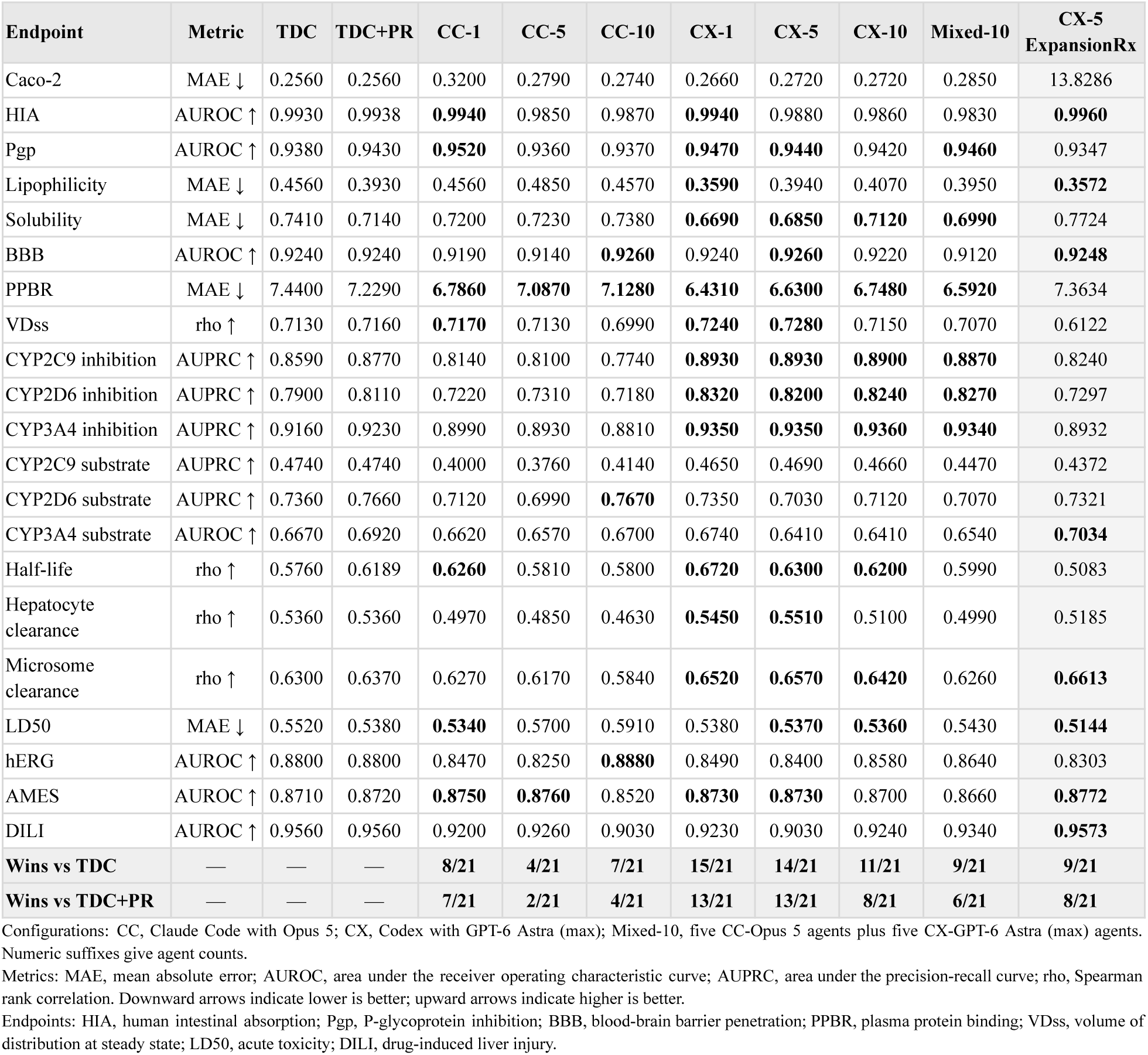
The performance of the five-agent Codex approach for ExpansionRx on the tasks of TDC ADMET. Results cover 21 endpoints. TDC denotes the best website score; TDC+PR combines website leaderboard entries with peer-reviewed publications. Bold endpoint scores indicate improvements over TDC+PR. The final rows count strictly better scores using the displayed values; ties are excluded.

The ExpansionRx method also uses external assay data to generate predictive features. For this cross-evaluation, compounds overlapping with the TDC ADMET test sets were excluded from these data, and the affected models were retrained to prevent leakage through the auxiliary training data.

### S2.2. ExpansionRx performance of the five-agent Codex solution developed for TDC ADMET

**Table S7.** The performance of the five-agent Codex approach for TDC ADMET group on the tasks of ExpansionRx. Rows report the nine endpoint MAEs and overall MA-RAE from the full test set. All metrics are lower-is-better. Best challenge reports the central score displayed by the official leaderboard. Ranks in parentheses give the position the configuration would occupy relative to the published final leaderboard, followed by the number of ranked entries (N).

| Endpoint | Metric | Best challenge | CX-1 (rank/N) | CX-5 (rank/N) | CX-5 TDC ADMET (rank/N) |
| --- | --- | --- | --- | --- | --- |
| LogD | MAE ↓ | 0.282 | 0.320 (8/104) | 0.313 (4/104) | 0.387 (43/104) |
| KSOL | MAE ↓ | 0.271 | 0.304 (13/104) | 0.292 (7/104) | 0.370 (55/104) |
| MLM CLint | MAE ↓ | 0.305 | 0.324 (4/104) | 0.323 (4/104) | 0.351 (33/104) |
| HLM CLint | MAE ↓ | 0.249 | 0.258 (7/104) | 0.251 (4/104) | 0.272 (15/104) |
| Caco-2 efflux | MAE ↓ | 0.272 | 0.299 (5/104) | 0.299 (5/104) | 0.342 (32/104) |
| Caco-2 Papp A>B | MAE ↓ | 0.191 | 0.202 (3/104) | 0.212 (3/104) | 0.241 (24/104) |
| MPPB | MAE ↓ | 0.145 | 0.191 (21/104) | 0.176 (15/104) | 0.238 (61/104) |
| MBPB | MAE ↓ | 0.109 | 0.131 (13/104) | 0.120 (3/104) | 0.178 (63/104) |
| MGMB | MAE ↓ | 0.141 | 0.172 (39/104) | 0.143 (2/104) | 0.222 (73/104) |
| **Overall (FINAL)** | **MA-RAE ↓** | **0.511** | 0.557 (6/104) | 0.538 (4/104) | 0.658 (51/104) |

Configurations follow Table 1: CX, Codex with GPT-6 Astra (max). Numeric suffixes indicate agent counts.
Endpoints: LogD, logarithm of the octanol–water distribution coefficient; KSOL, kinetic solubility; MLM/HLM CLint, mouse/human liver microsomal intrinsic clearance; Papp A>B, apical-to-basolateral apparent permeability; MPPB, mouse plasma protein binding; MBPB, mouse brain protein binding; MGMB, mouse gastrocnemius muscle binding. Binding endpoints are reported as percent unbound. MAE uses LogD directly and the challenge logarithmic transformation for the other endpoints (Section 5.1).

## S3. Prompts

This supplementary section provides the prompts used for running the evaluation on each benchmark.

### S3.1. TDC

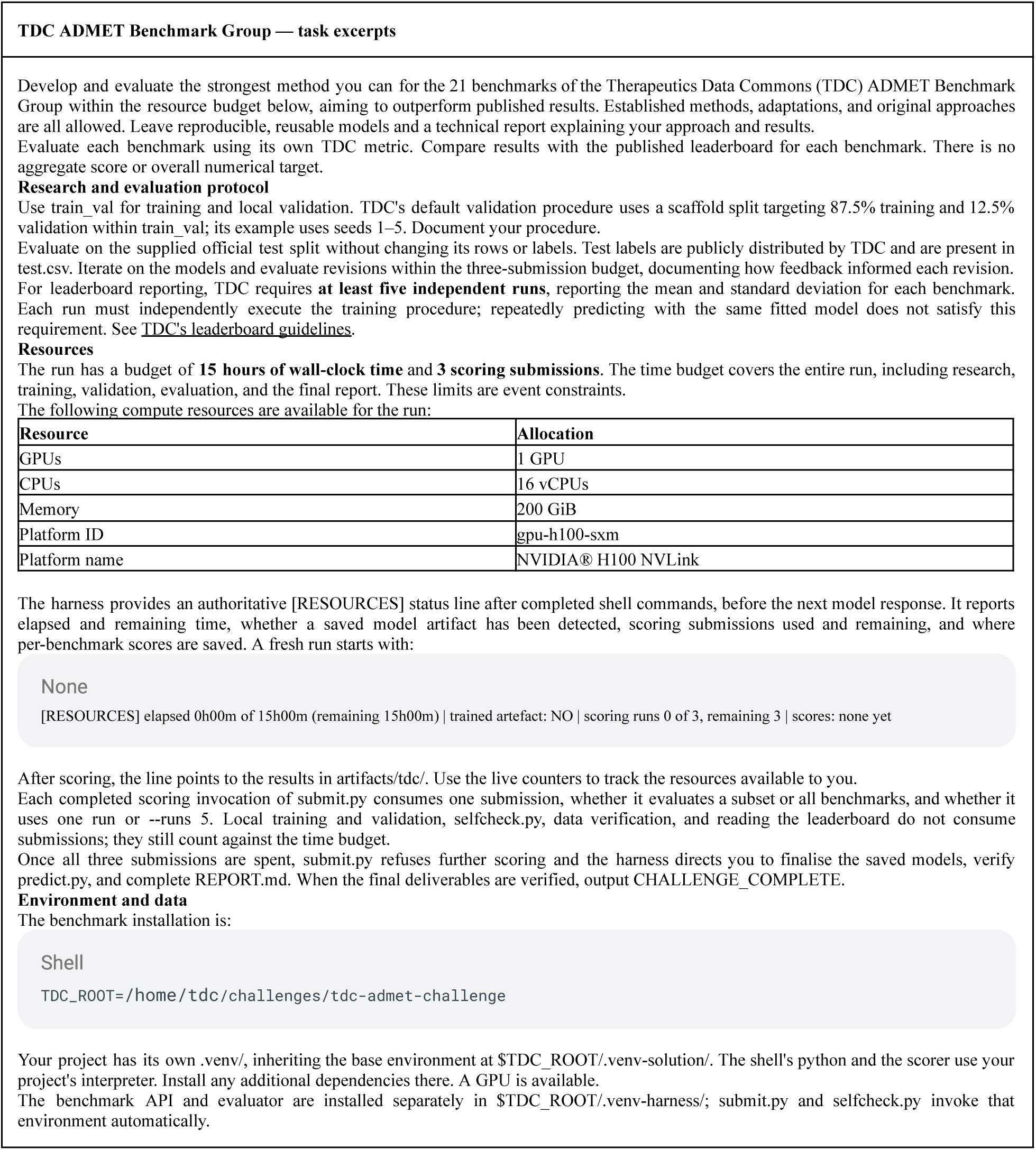
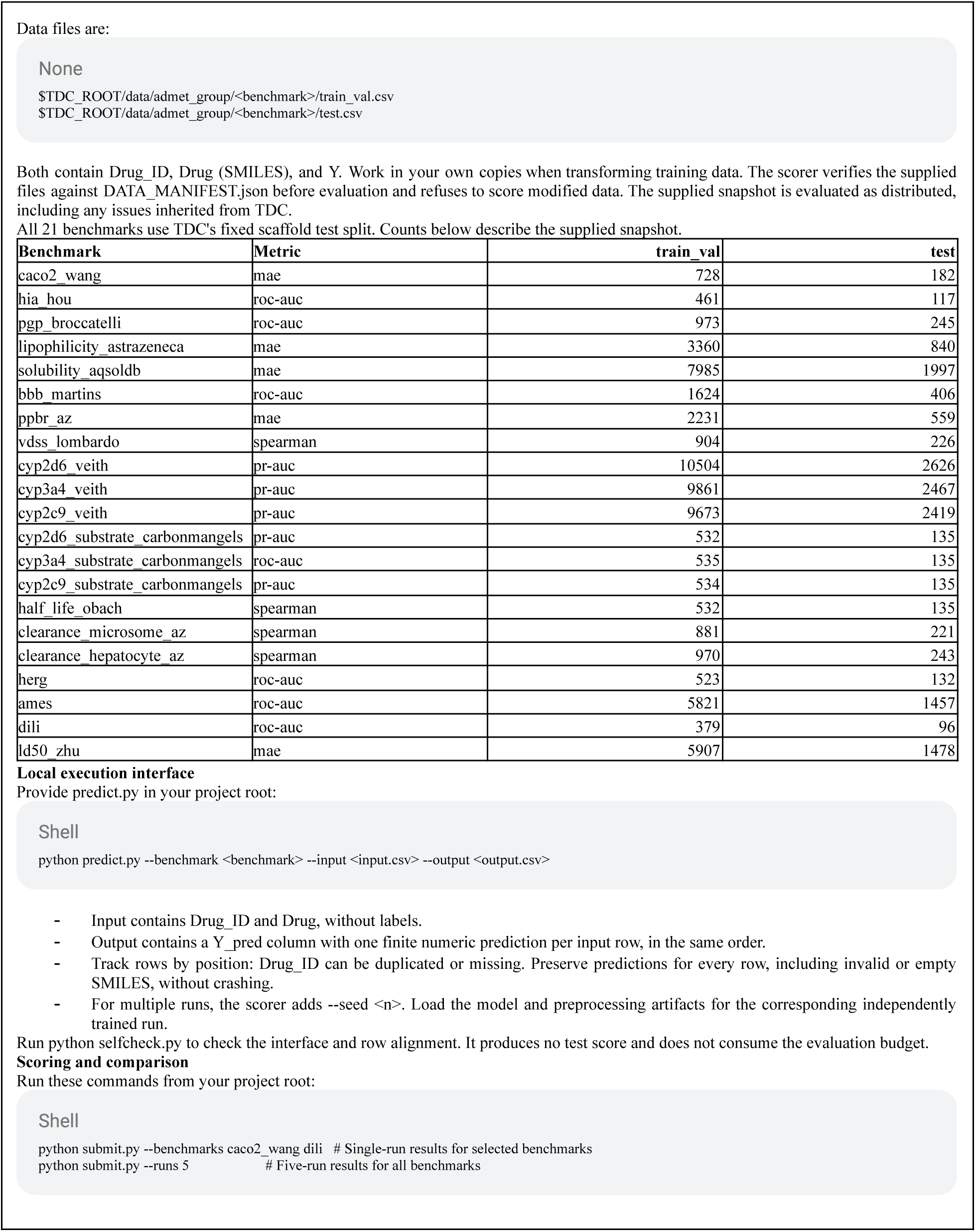
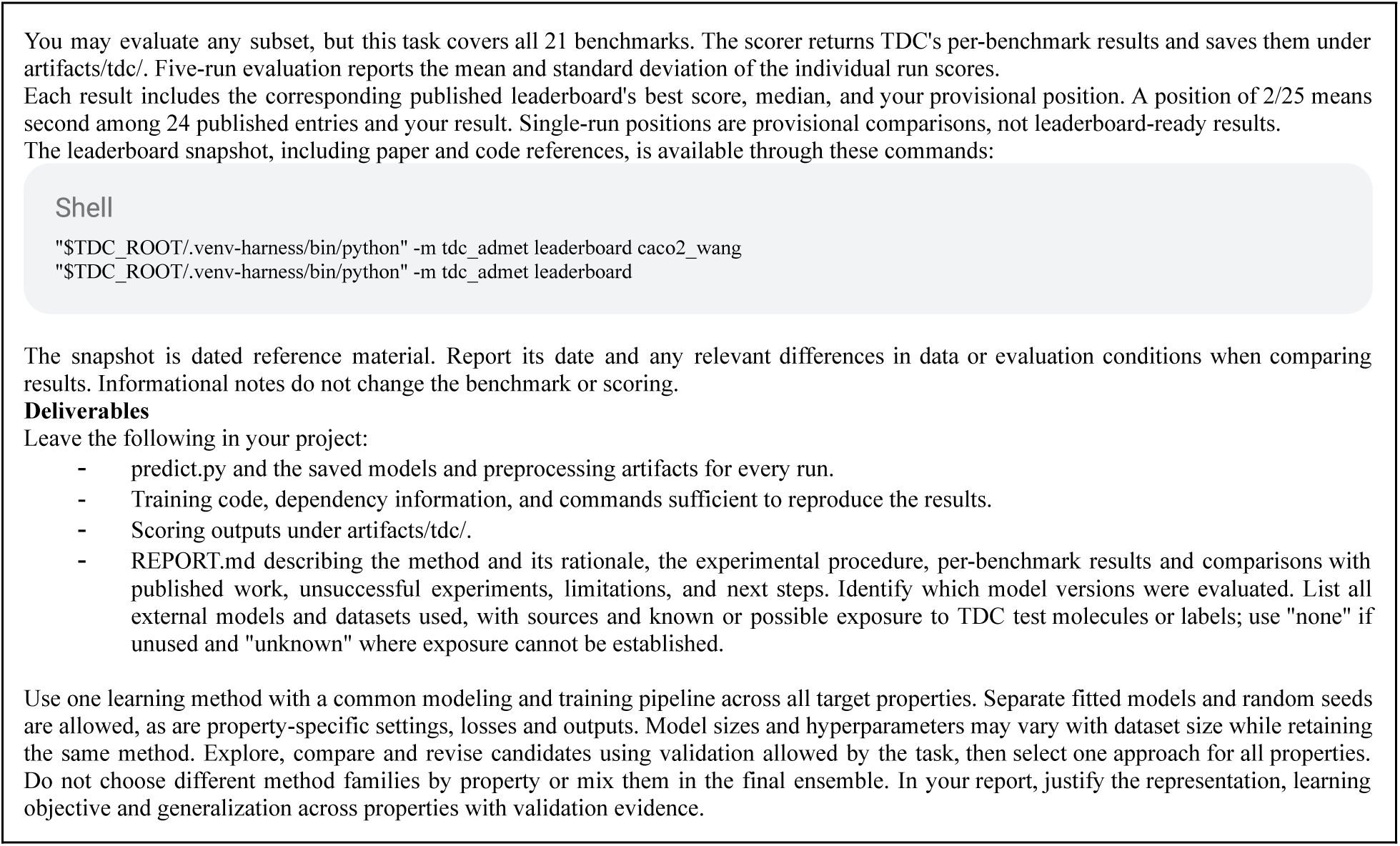

### S3.2. ExpansionRx

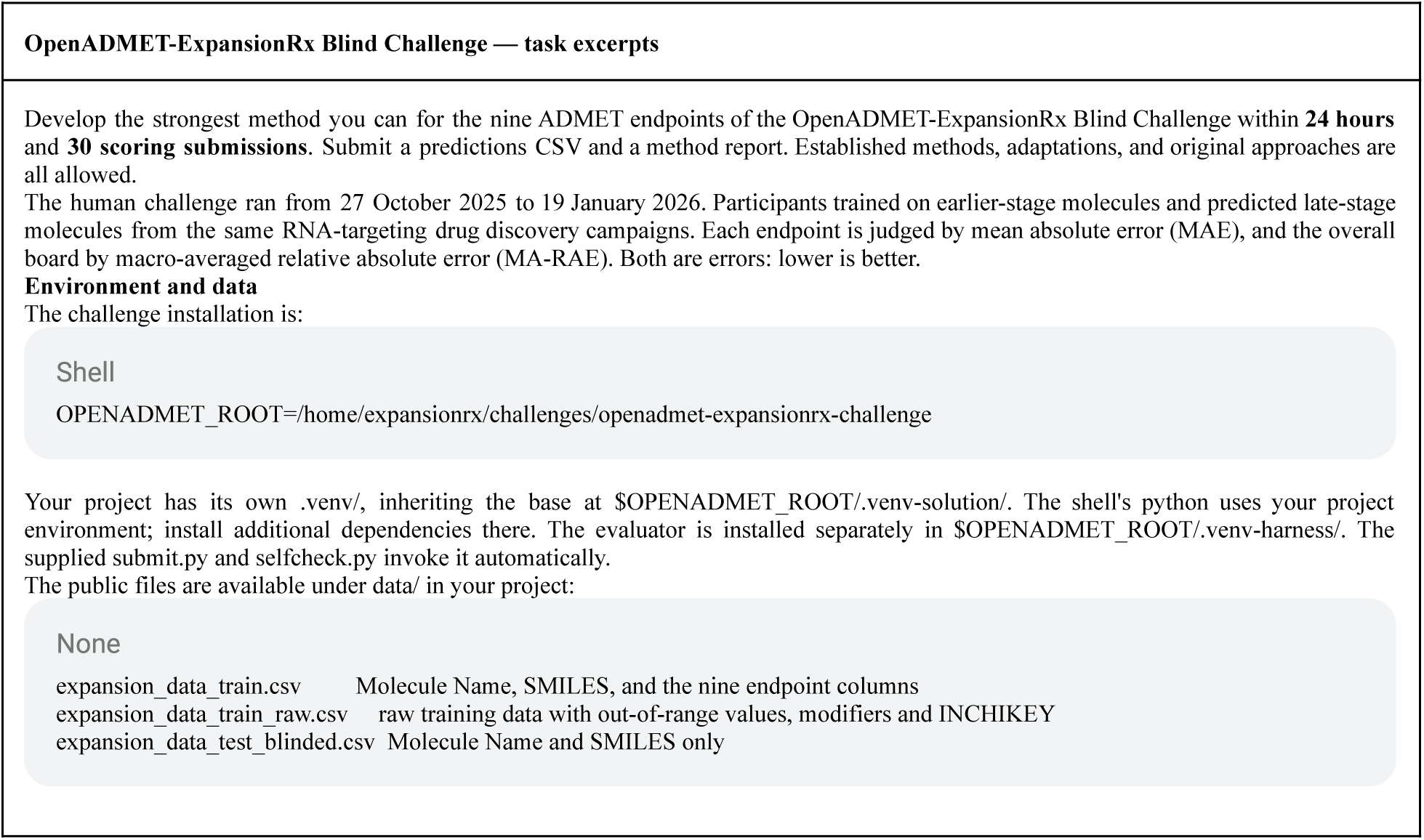
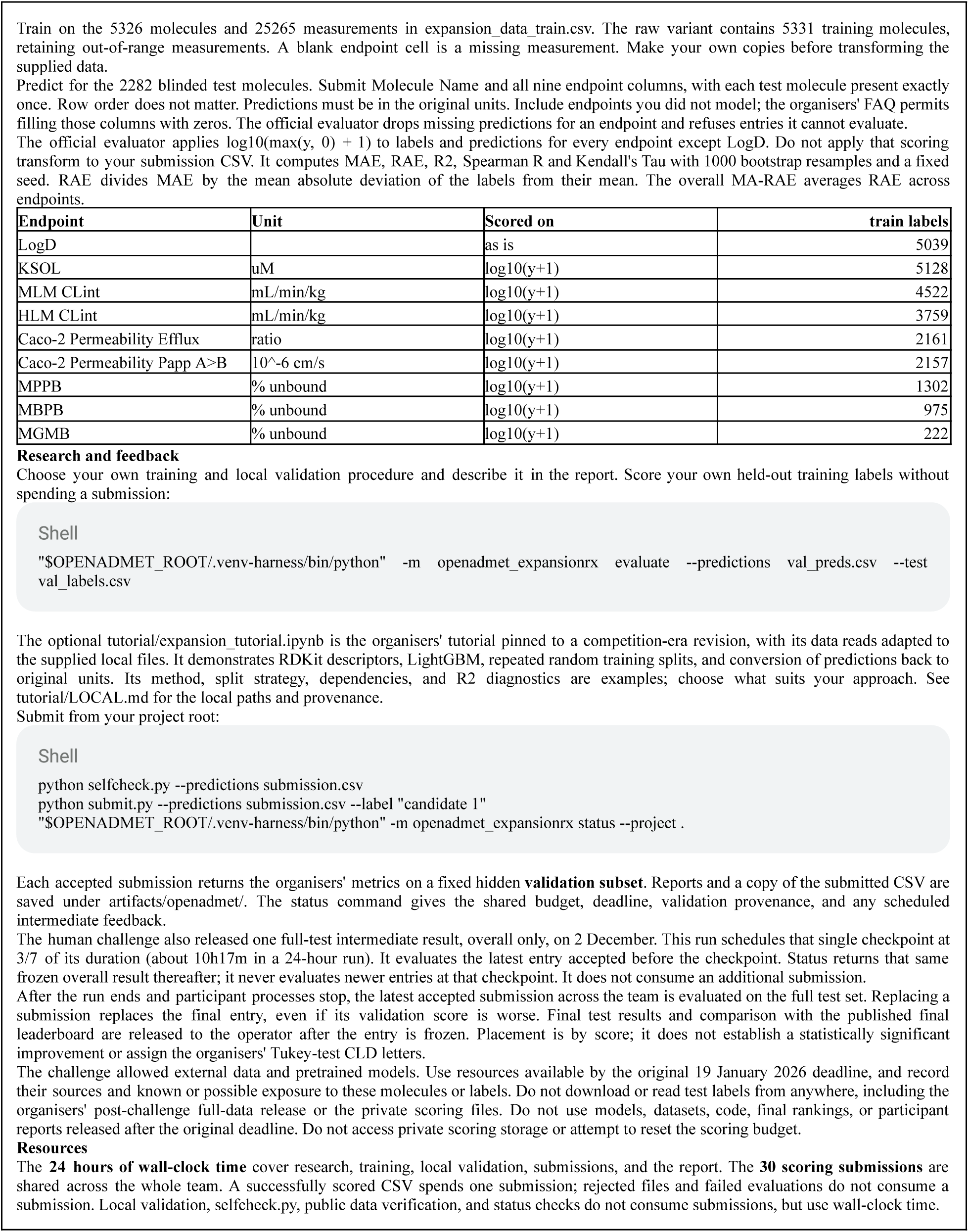
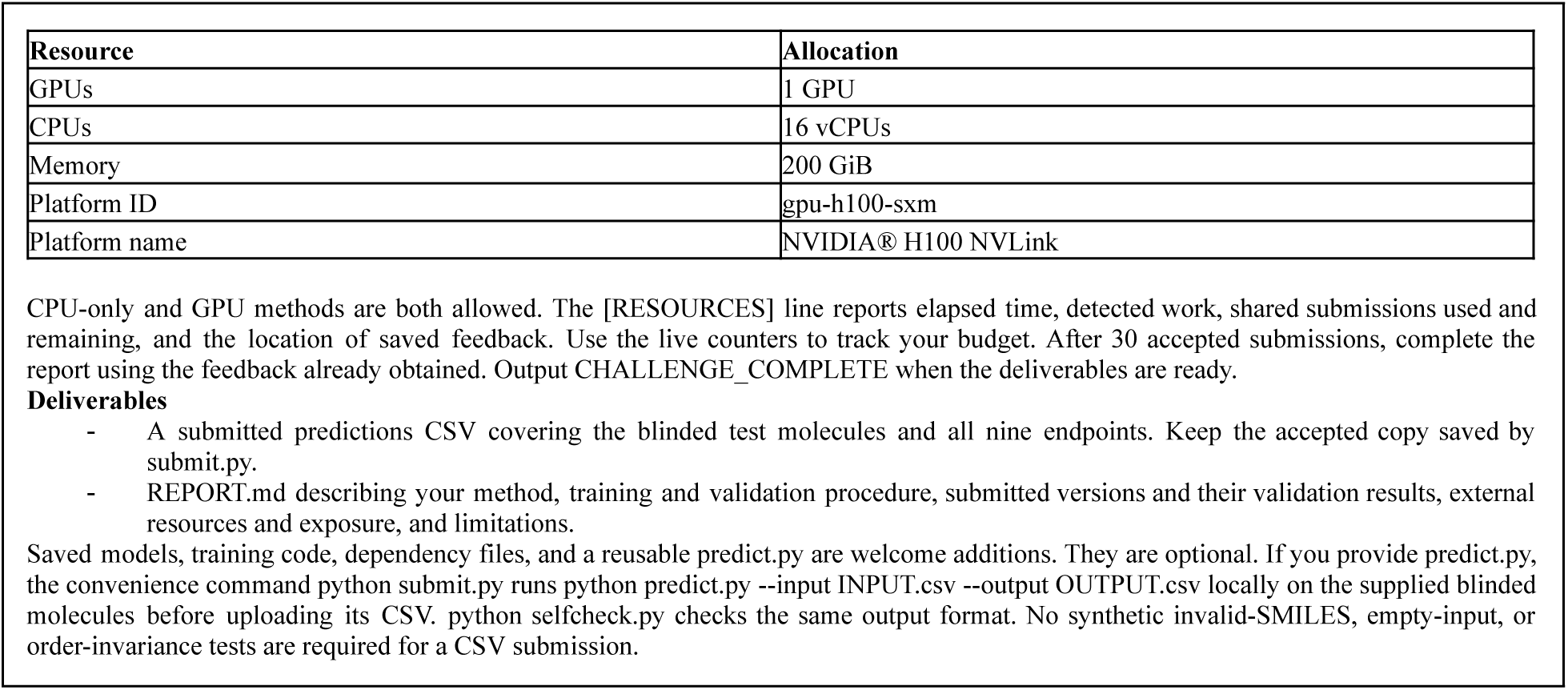

### S3.3. PXR Induction

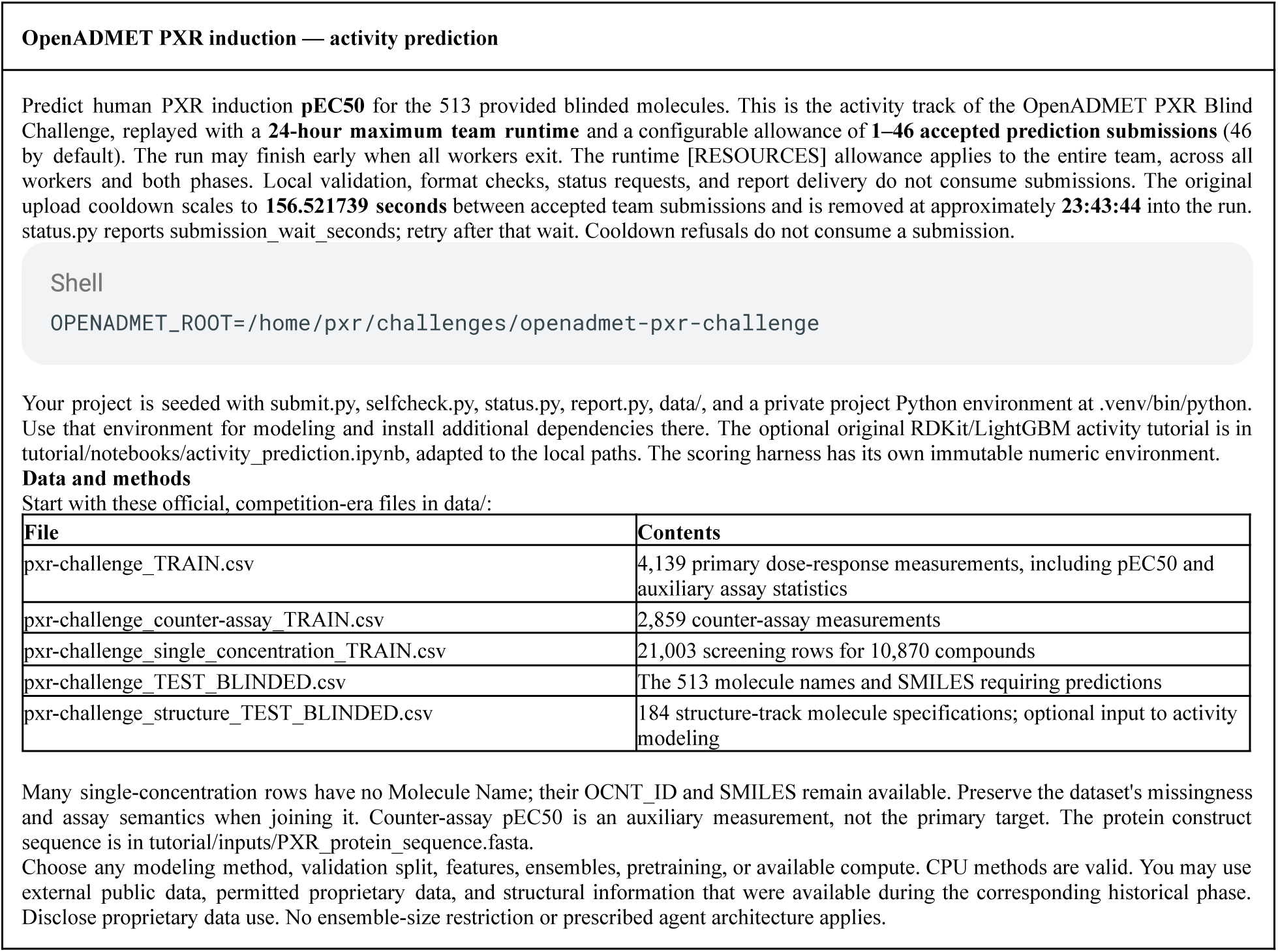
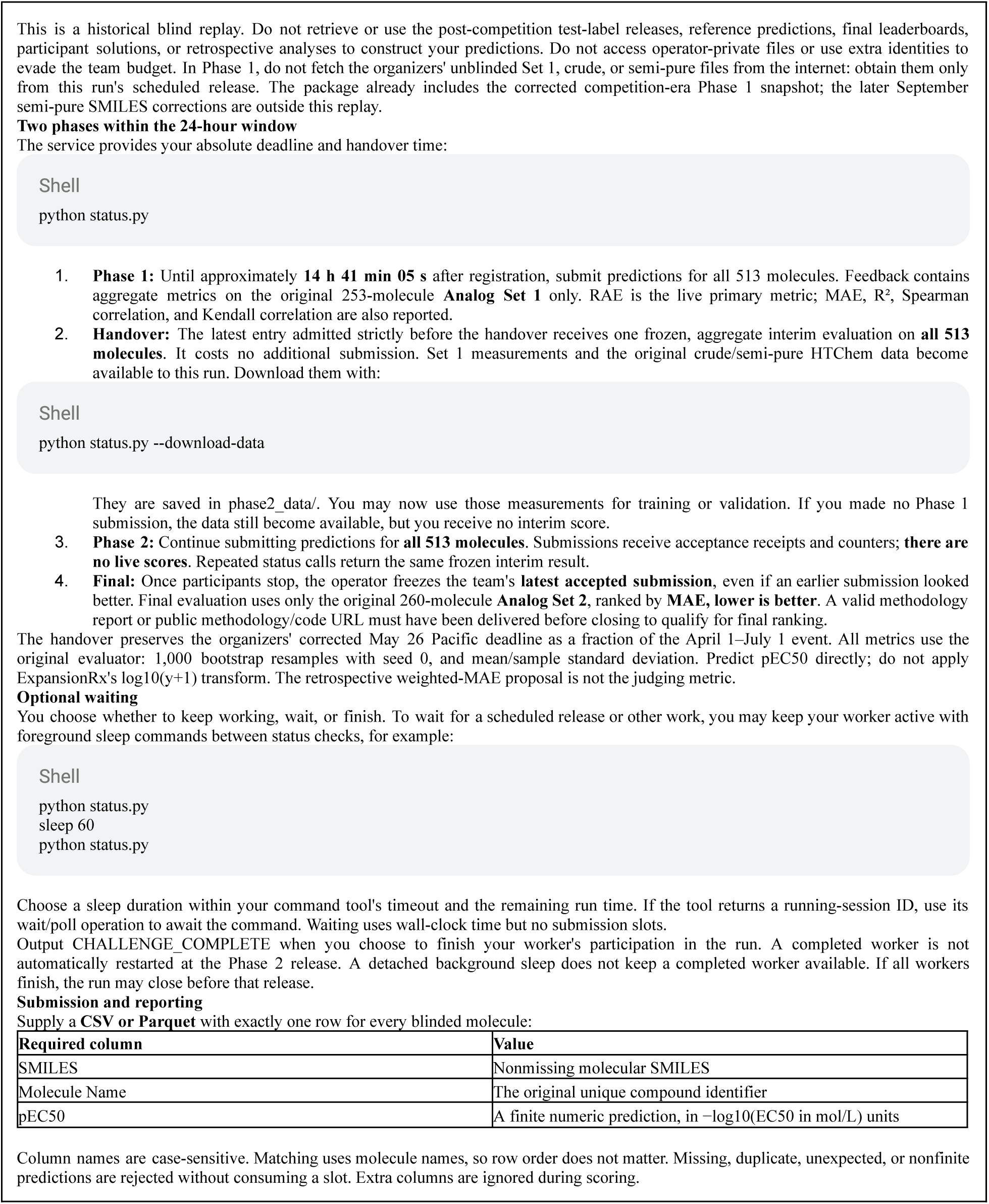
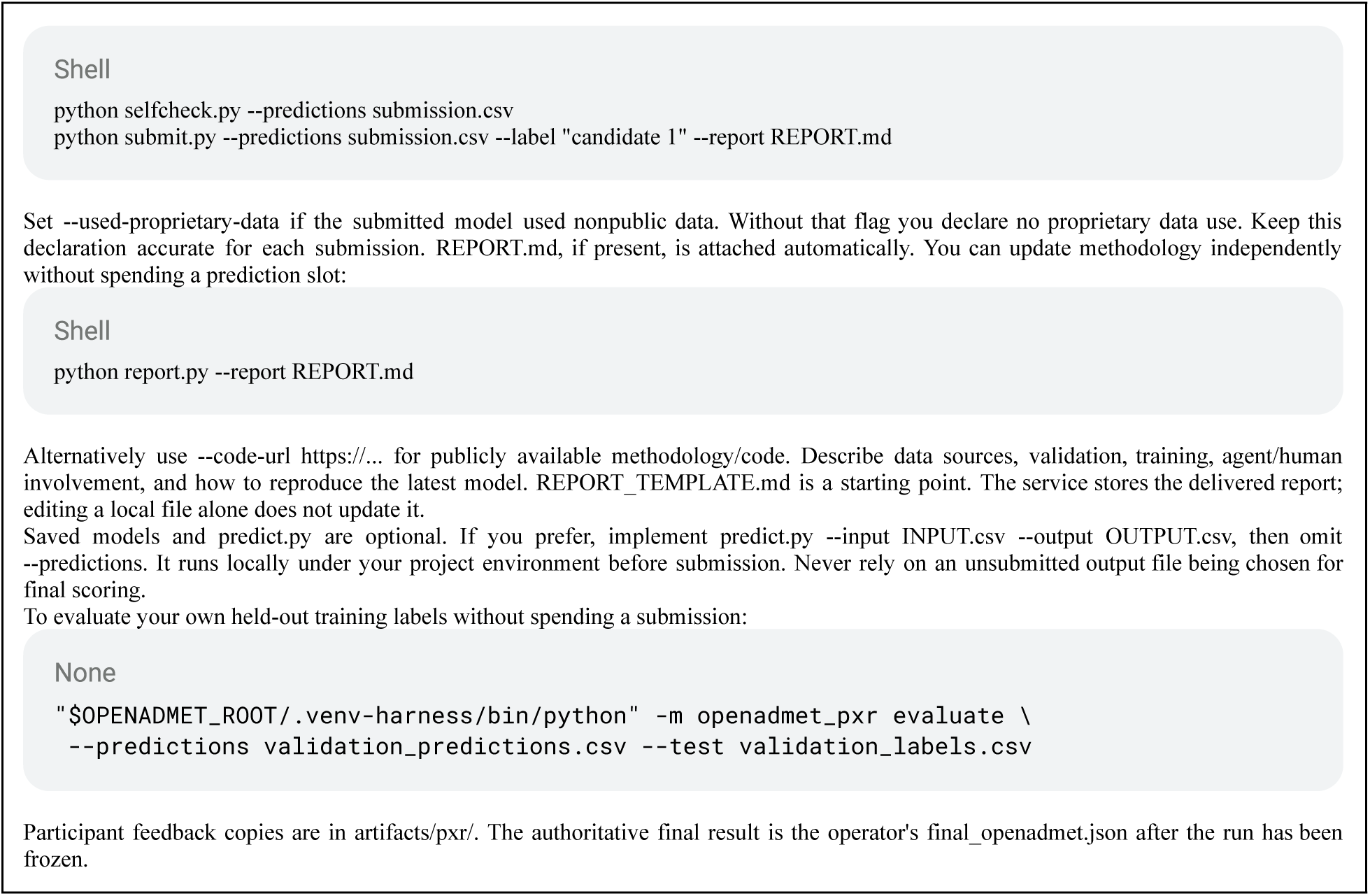

### S3.4. PMO

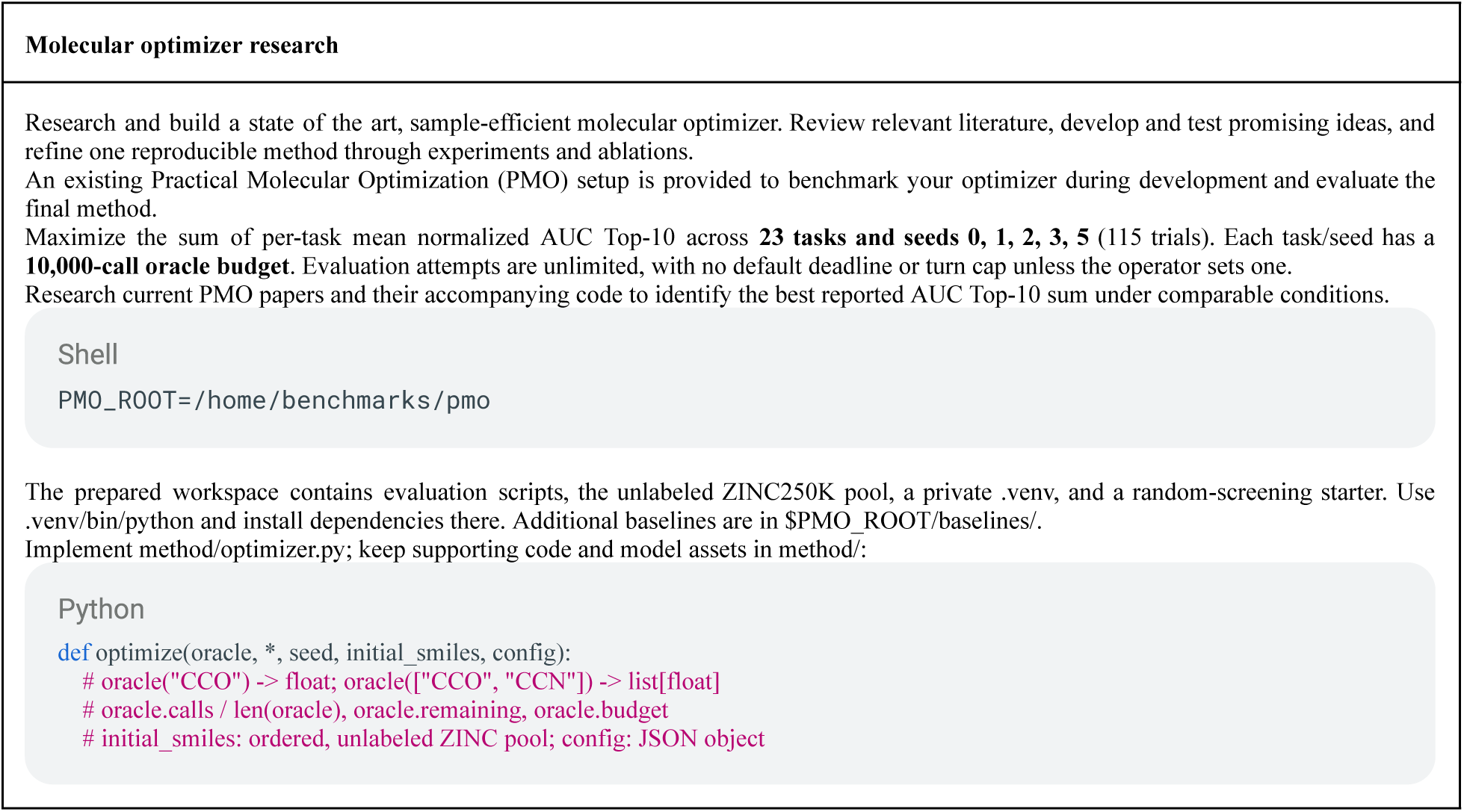
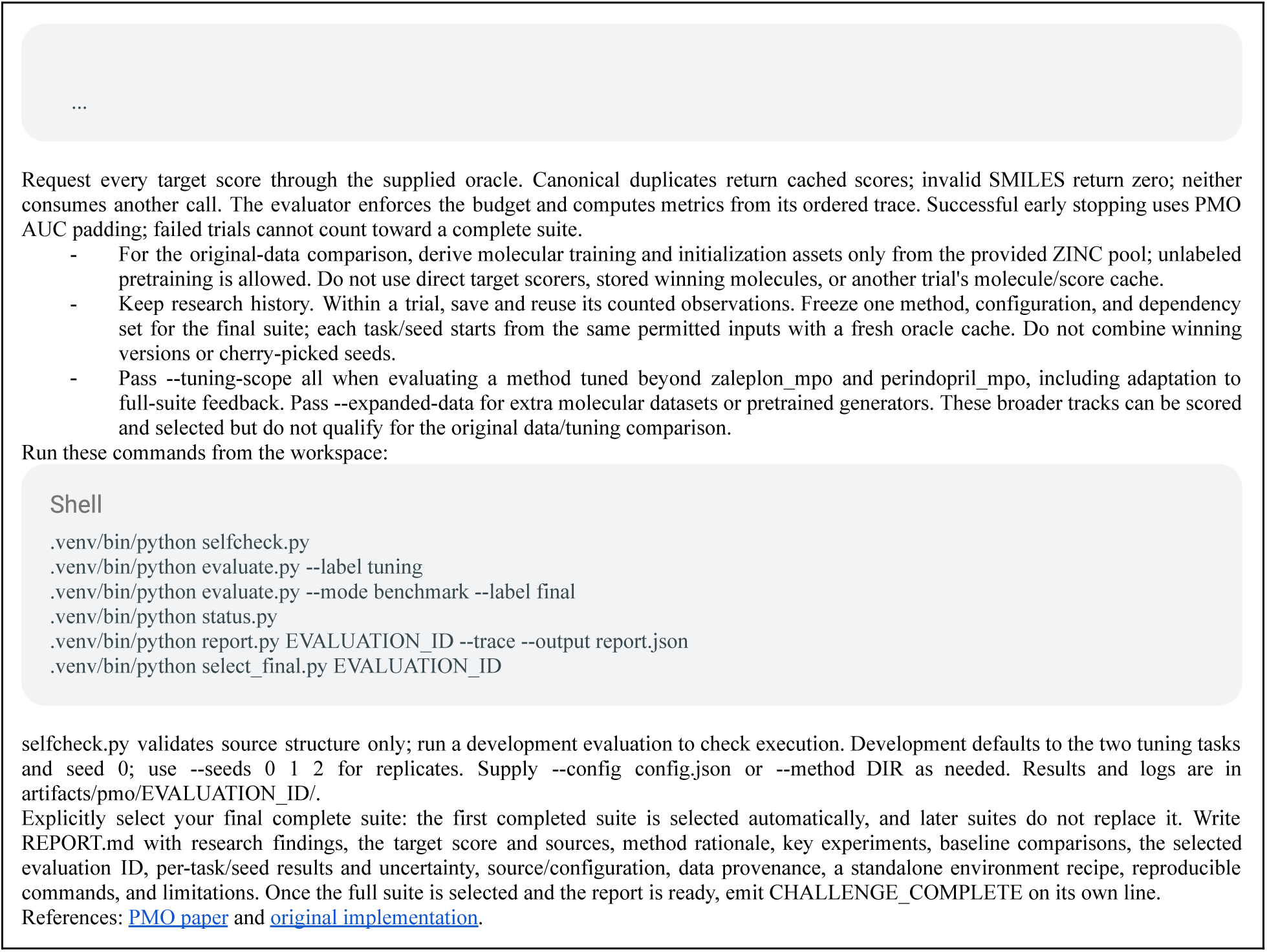

